# Dissecting and directing pathology foundation models

**DOI:** 10.64898/2026.06.12.731496

**Authors:** Chanwoo Kim, Jakub Kaczmarzyk, Deepika Savant, Zhen Zhao, Peter K Koo, Su-In Lee

## Abstract

Foundation models (FMs) are central to digital pathology, encoding histology images into dense *embeddings* for facilitating diagnostic classification, molecular alteration prediction, and clinical outcome modeling. However, the opacity of these embeddings renders FM-based systems “black boxes,” limiting their trustworthiness for clinical translation and utility for scientific discovery. Here, we introduce PICASSO (Pathology Image Concept Atlas built via SparSe dictiOnary learning), a framework that makes pathology FMs interpretable and controllable. PICASSO decomposes FM embeddings into human-interpretable visual *concepts* using a sparse autoencoder. It is trained on more than 120 million tissue patches across 32 cancer types, producing the first pan-cancer atlas of histomorphological concepts. We demonstrate that PICASSO enables diverse downstream applications of FM embeddings by exposing interpretable structure within learned representations and supporting concept-level intervention. It enables auditing of clinical model behavior by revealing the morphological features driving predictions. Beyond transparency and validation, PICASSO enables the discovery of new biological insights; for example, it identified hobnailing epithelial morphology as a previously unrecognized biomarker of EGFR mutations in lung adenocarcinoma. By linking PICASSO-derived concepts with spatial transcriptomics, we uncover associations between morphological patterns and gene expression programs. Furthermore, PICASSO allows suppression of concepts associated with technical artifacts, thereby reducing model reliance on spurious signals. Finally, PICASSO enables controlled manipulation of learned concepts to generate counterfactual embeddings for exploratory therapeutic analysis, such as modulating tumour-infiltrating lymphocyte density to assess impacts on predict survival outcomes. Together, PICASSO provides a principled framework for transforming pathology FMs into platforms for mechanistic insight and discovery.

## Introduction

Histopathology slides, now increasingly digitized, are a clinically indispensable modality routinely collected across medicine for diagnosis and therapeutic guidance. Pathologists examine these specimens to determine tumor presence and subtype, forming the basis of many downstream clinical decisions. Over the past decade, digital pathology has rapidly evolved into a transformative field, with artificial intelligence (AI) achieving strong performance in routine diagnostic tasks and showing promise in reducing pathologists’ workload^1–3^.

A central driver of this progress is the emergence of pathology foundation models (FMs), trained in an unsupervised manner on vast collections of histology images to learn rich representations of tissue architecture and cellular morphology^4–7^. These *embeddings*, encoded as dense numerical vectors, can be input to task-specific models performing standard pathology tasks (Fig. 1a)—such as cancer diagnosis including subclassification and grading—or more challenging tasks beyond routine human assessment, including inferring molecular alterations and predicting survival. Remarkably, these capabilities are achieved directly from routine hematoxylin and eosin (H&E) images, bypassing specialized assays. For example, a recent prospective study showed that a pathology FM could accurately predict epidermal growth factor receptor (*EGFR*) mutations from H&E slides in lung adenocarcinoma, improving specimen utilization, cost efficiency, turnaround time, and diagnostic insight^8^.

**Fig. 1.**
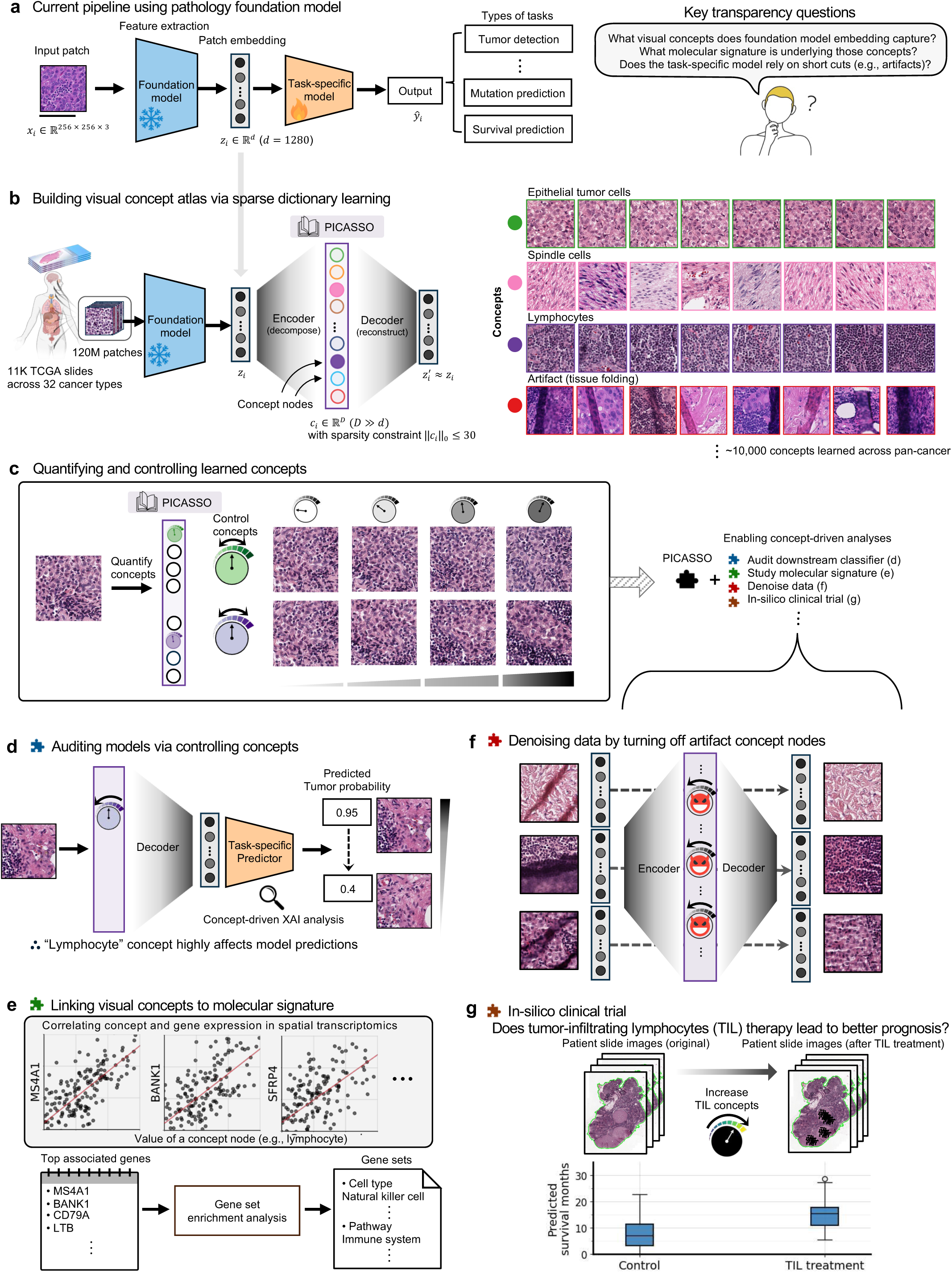
Overview of PICASSO framework and its usage examples. See next page for caption. a, Current pipeline using a pathology foundation model. A foundation model encodes histopathology image patches into high-dimensional embeddings, which are used by task-specific models to perform downstream clinical tasks. Scale bar, 100 µm. b, Building a visual concept atlas from large-scale pan-cancer data. Trained on over 120 million histology patches from over 11,000 TCGA slides spanning 32 cancer types, PICASSO learns the visual vocabulary underlying these embeddings. PICASSO decomposes dense FM embeddings into a sparse set of interpretable ‘concept nodes,’ each representing specific histological features. There are around 10,000 concepts, of which the activation frequencies are higher than 0.1%. (Supplementary Fig. 3). c, Quantifying and controlling concepts. Individual concept activations can be selectively increased or decreased, enabling direct semantic control of FM embeddings and concept-driven analyses. d, Auditing downstream models via concept control. Manipulating specific concepts (e.g., lymphocytes) and observing the resulting changes in model outputs (e.g., tumor probability) reveals the morphological drivers of predictions. e, Linking visual concepts to molecular signatures. Concept activations are correlated with gene expression measurements from spatial transcriptomics data to identify underlying gene expression programs. f, Denoising data by suppressing artifact concept nodes. Turning off artifact-related concept nodes (e.g., tissue folding) cleans embeddings of non-histopathological features and mitigates downstream models’ reliance on spurious signals. g, In-silico clinical trials via concept-level intervention. PICASSO can manipulate clinically meaningful concepts in pathology images, enabling the simulation of therapeutic interventions, such as increasing the tumor-infiltrating lymphocyte (TIL) in silico to assess predicted therapy responsiveness.

Despite these advances, FM-based systems—AI systems that use FM embeddings to perform downstream tasks— remain essentially “black boxes,” posing major obstacles to clinical deployment and raising concerns about safety, trust, and regulatory approval^9^. These limitations arise from three complementary gaps: (i) limited interpretability of FM embeddings, (ii) lack of biological grounding of learned representations, and (iii) susceptibility to spurious or non-biological features in model predictions (Fig. 1a). First, FM embeddings constitute the core representation in these systems, yet the visual information they encode remains poorly understood. These embeddings are dense numerical vectors of approximately 1,000-2,000 dimensions that capture a wide spectrum of visual concepts, from morphologic features and cellular components to tissue organization. However, individual dimensions lack clear semantic interpretation^10,11^, limiting the ability to query or control specific visual concepts encoded in these representations. Second, although FM-based systems perform tasks beyond routine human assessment, such as predicting prognosis or molecular alterations directly from H&E images, the morphological cues driving these predictions are not well linked to underlying molecular mechanisms. Establishing these links—for example, by anchoring histomorphology to gene expression programs—could reveal gene regulatory programs underlying morphological biomarkers and provide mechanistic insight into disease. Third, safe clinical deployment requires ensuring that model predictions are driven by biologically meaningful histopathologic features rather than artifacts such as tissue folds or slide preparation variability.

To address these critical challenges, we present PICASSO (Pathology Image Concept Atlas built via SparSe dictiOnary learning), a unified framework that converts opaque embeddings into human-interpretable visual concepts and enables explicit control over those concepts (Fig. 1b-c). The core challenge is that FM embeddings are dense and entangled, with individual dimensions encoding multiple concepts, a phenomenon known as polysemanticity^10,12,13^. We address this using a sparse autoencoder (SAE) grounded in dictionary learning (see Methods), which decomposes each embedding into a sparse linear combination of learned basis vectors corresponding to interpretable visual concepts. PICASSO learns a large *dictionary* of visual concepts captured by existing pathology FMs across 32 cancer types. Each dictionary entry defines an interpretable ‘concept node’ representing specific histological features—such as lymphocytes, stromal organization, or nuclear pleomorphism—that can be directly visualized and quantified (Fig. 1b). Trained on all available diagnostic H&E slides from TCGA—over 120 million patches from more than 11,000 slides spanning 32 cancer types—PICASSO constitutes, to our knowledge, the first pan-cancer atlas of visual concepts learned at scale from pathology FMs.

While the SAE reveals concepts encoded in FM embeddings, interpreting these concepts remains challenging, particularly in histopathology, where images are complex and diagnostically relevant differences are often subtle. Prior work in other domains has typically interpreted SAE concepts by inspecting high-activation exemplars^10^. However, in histopathology, such exemplars are often insufficient, as many clinically meaningful features are better understood through contrastive visual comparison^14–17^. To address this, we developed Emb2Img, a complementary diffusion model that reconstructs realistic histology patches from FM embeddings (see Methods). By systematically modulating individual concept nodes and visualizing the resulting morphological changes (Fig. 1c), Emb2Img enables interpretation through controlled perturbations rather than static exemplars, analogous to counterfactual image-based explanations in explainable AI^14,16^. This framework enables a *human-in-the-loop* workflow in which pathologists evaluate these perturbations and anchor each concept to histopathologically meaningful morphology. Through close collaboration with expert pathologists during concept verification, we confirmed that the learned concepts correspond to recognizable and clinically relevant histological features.

We demonstrate how PICASSO enables transparent use of FMs across diverse applications (Fig. 1d–g). First, PICASSO enables systematic auditing of downstream models by modulating individual concepts to observe resulting shifts in model outputs, thereby revealing the morphological features driving predictions (Fig. 1d). Second, PICASSO grounds these insights in molecular biology by linking histological concepts to genes and pathways through spatial transcriptomics (Fig. 1e). Third, concept-level modulation improves model robustness by suppressing artifact-related concepts, such as tissue folds, thereby reducing reliance on spurious cues (Fig. 1f). Fourth, concept manipulation enables systematic virtual experiments through counterfactual perturbations, such as simulating tumour-infiltrating lymphocyte therapy to probe therapeutic efficacy (Fig. 1g).

Taken together, PICASSO provides a principled framework for transforming pathology FMs into interpretable, controllable platforms for mechanistic insight, biological discovery, and clinically relevant hypothesis generation.

## Results

### Decomposing pathology FM embeddings into interpretable histopathological concepts

PICASSO translates opaque pathology FM embeddings into human-interpretable visual concepts using an SAE (Fig. 1b). The SAE maps each dense embedding into a high-dimensional concept space, decomposing it into interpretable morphological features, where each dimension represents a *concept activation* quantifying the presence of a specific concept within the image. Sparsity constrains only a small subset of concepts to be active for each image while preserving information encoded in the original embedding. Because PICASSO operates on FM embeddings, our framework generalizes across pathology FMs. Here, we applied PICASSO to embeddings from Virchow2^4,18^, a high-performing pathology FM with consistently strong performance across downstream clinical tasks (Supplementary Figs. 1-2).

As an initial validation of the PICASSO framework, we evaluated its decomposition according to two criteria: *fidelity*, defined as the extent to which extracted concepts preserve information from the original embeddings, and *interpretability*, defined as the degree to which these concepts correspond to human-recognizable histological features. We first assessed fidelity by measuring the proportion of variance in the original embeddings explained by reconstructions derived from PICASSO concept activations (Fig. 2a; Methods). We benchmarked PICASSO against classical unsupervised embedding decomposition approaches, including principal component analysis (PCA) and *k*-means clustering, both of which can be viewed as dictionary learning^19^. In PICASSO, the sparsity hyperparameter *k* controls the number of active concept nodes and thereby the granularity of the learned representations. Across varying numbers of components (*k* for PICASSO, the number of principal components for PCA, and the number of clusters for *k*-means), PICASSO consistently achieved the highest fidelity. Notably, with only *k* = 30 active concepts, PICASSO explained over 80% of the variance in Virchow2 embeddings, indicating that FM embeddings can be compactly represented using a relatively small number of concepts. We therefore used *k* = 30 in all subsequent analyses.

**Fig. 2.**
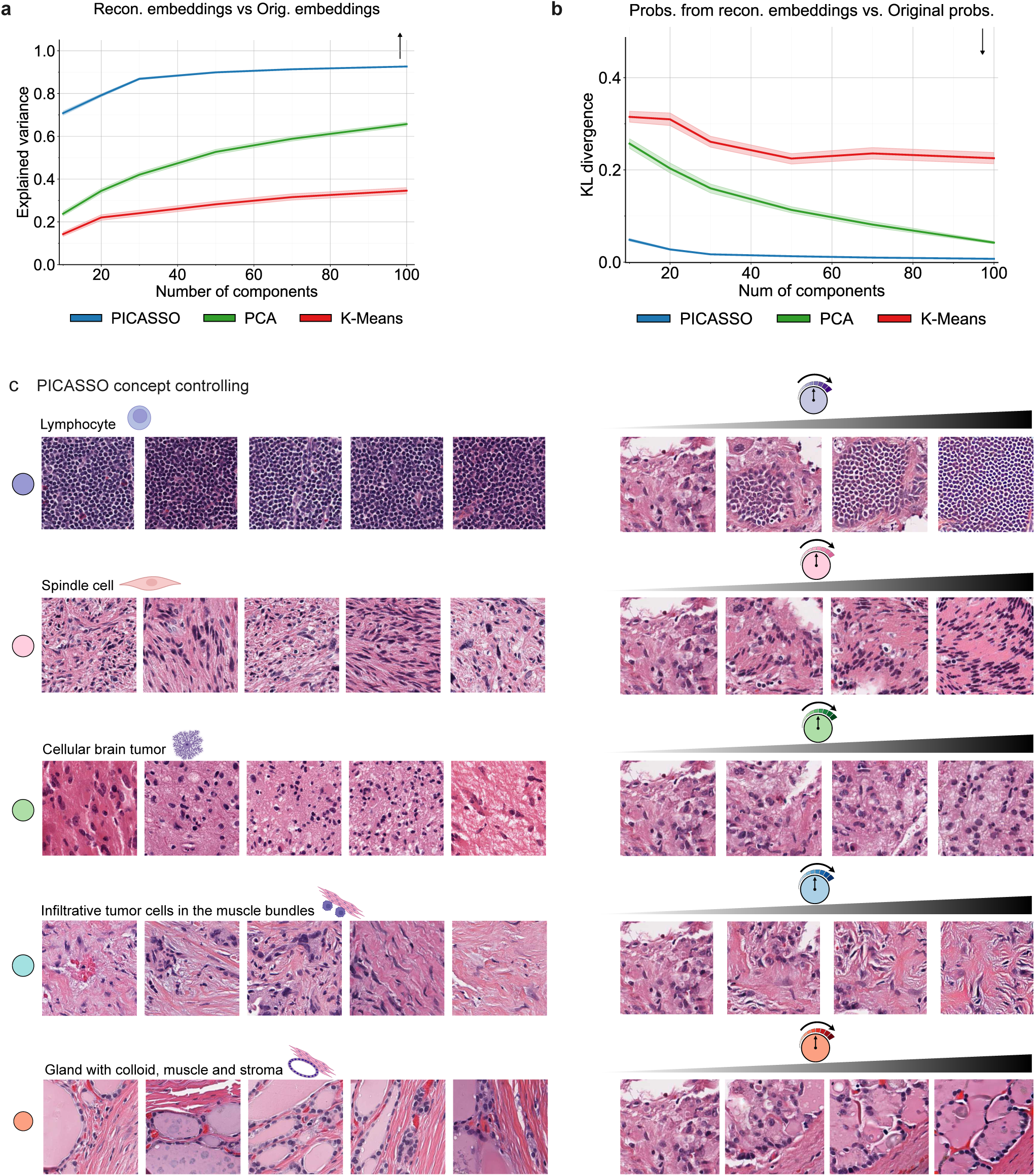
Fidelity and interpretability of PICASSO concepts. a, Fidelity at the embedding level. The y-axis shows the proportion of variance in original FM embeddings explained by reconstructed embeddings, evaluated across different numbers of components to represent embeddings. Higher values are better (indicated by arrow). The center line represents the mean explained variance calculated over *n* = 10, 000 TCGA test samples, with the shaded area indicating the 95% confidence interval. b, Fidelity at the downstream prediction level. The y-axis shows the Kullback–Leibler (KL) divergence between predictive probabilities produced from reconstructed embeddings and those from the original embeddings. Lower values are better (indicated by arrow), indicating less deviation. The center line represents the mean KL divergence calculated over *n* = 100, 000 downstream test samples, with the shaded area indicating the 95% confidence interval. c, Examples of concepts learned by PICASSO and their morphological interpretations. For each concept, the left panel shows the real image examples that most strongly activate the corresponding concept node. The right panel shows synthetic images generated by Emb2Img as the activation of that concept is progressively increased, illustrating semantic control over histological features.

This fidelity extended to downstream clinical tasks. Predictive outputs across diverse models remained highly consistent when original embeddings were replaced by reconstructed embeddings (Kullback–Leibler divergence <0.01; Fig. 2b), confirming that PICASSO preserves information essential for FM-based prediction.

We next evaluated the histologic interpretability of the learned concepts in collaboration with expert pathologists. As an initial observational analysis, we examined image patches that maximally activated individual concept nodes. In some cases, these exemplars revealed coherent and clinically recognizable morphologic patterns, including dense sheets of lymphocytes or spindle cells, and glandular structures with colloid content, among others (Fig. 2c). However, exemplar-based interpretation alone is often insufficient in complex histology.

To directly characterize the tissue morphology represented by individual concepts, we complemented this observational analysis with a perturbation-based approach using Emb2Img, a diffusion model that reconstructs realistic histology patches from FM embeddings (Supplementary Fig. 4–6). By systematically increasing activation of individual PICASSO concepts while holding others fixed, and reconstructing the image patches with Emb2Img, we generated counterfactual image series that revealed how modulation of a single concept progressively altered tissue morphology (see Methods). For example, progressively increasing activation of a “lymphocyte” concept produced corresponding increases in lymphocyte density within reconstructed counterfactual images (Fig. 2c, right panels), enabling pathologists to isolate relationships between individual concepts and tissue morphology within complex histologic context. Across both exemplar-based observations and counterfactual perturbations, expert pathologists confirmed that the learned concepts correspond to recognizable and clinically meaningful histologic features.

In summary, these results demonstrate that PICASSO provides a high-fidelity, interpretable decomposition of pathology FM embeddings, establishing a foundation for systematic interpretation and control of pathology FMs.

### Constructing a pan-cancer atlas of morphological concepts

Having established that individual PICASSO concepts are faithful and interpretable, we next asked whether they collectively capture meaningful cellular and architectural features across cancers. Because PICASSO is trained at pan-cancer scale, it learns a broad morphological vocabulary spanning diverse tumor types. To characterize this landscape, we profiled concept activations across 6,812 TCGA whole-slide images from 14 cancer types and summarized the dominant morphological patterns within each cancer (Methods). The resulting atlas revealed distinct morphological “fingerprints” across malignancies (Fig. 3).

**Fig. 3.**
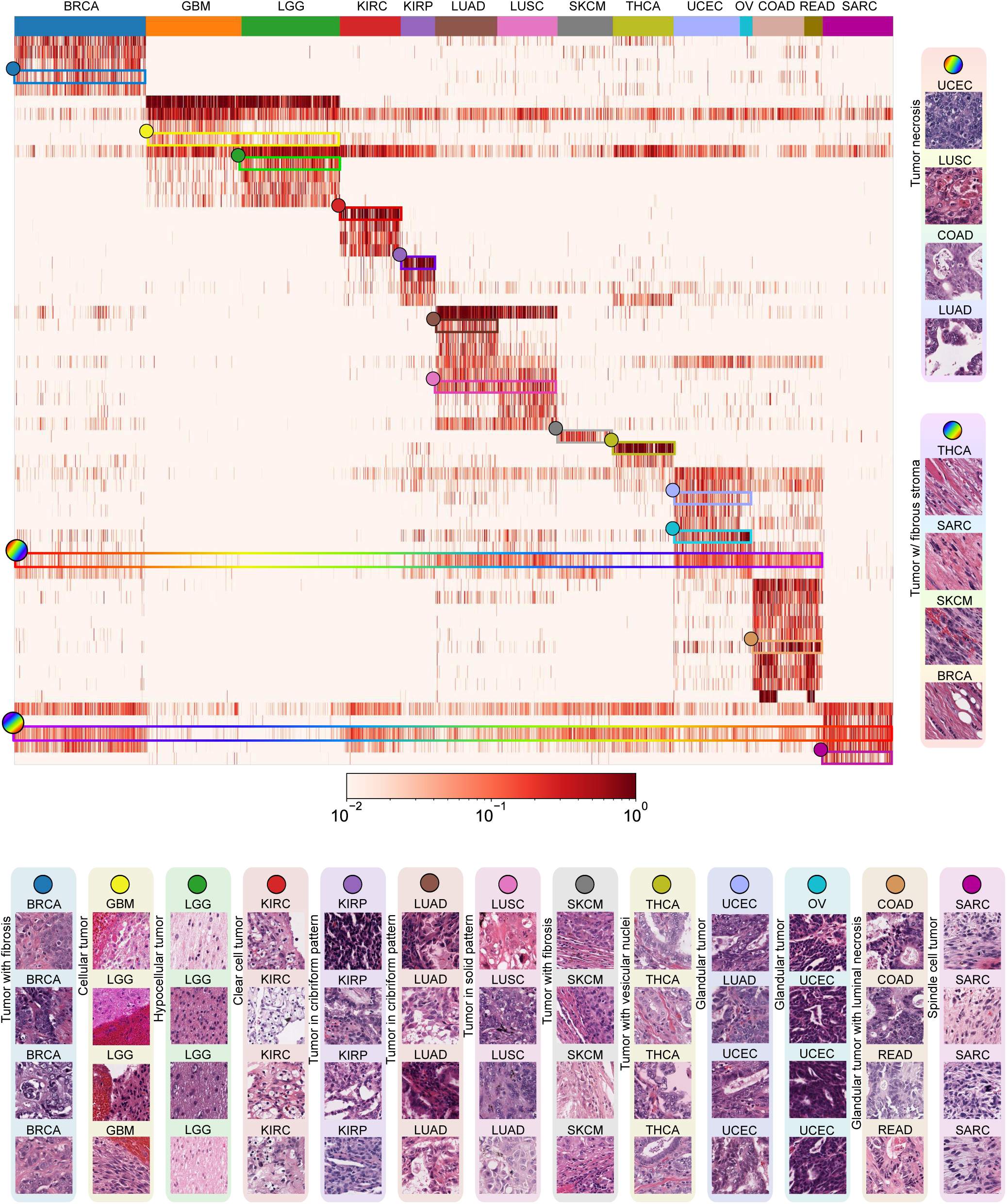
Pan-cancer heatmap of concept expression across 14 TCGA cancer types. Each column represents a whole-slide image (WSI) from an individual patient, grouped and color-coded by cancer type as indicated at the top. Each row corresponds to a distinct morphological concept learned by PICASSO. The color intensity represents the mean activation of each concept within the tumor regions of a WSI, plotted on a logarithmic scale. Higher intensity (darker color) signifies a more prominent presence of that morphological feature in the patient. The concepts (rows) are selected for their high variance across cancer types and are grouped by the cancer type in which they are most predominantly expressed. BRCA, breast invasive carcinoma; GBM, glioblastoma multiforme; LGG, low grade glioma; KIRC, kidney renal clear cell carcinoma; KIRP, kidney renal papillary cell carcinoma; LUAD, lung adenocarcinoma; LUSC, lung squamous cell carcinoma; SKCM, skin cutaneous melanoma; THCA, thyroid carcinoma; UCEC, uterine corpus endometrial carcinoma; OV, ovarian serous cystadenocarcinoma; COAD, colon adenocarcinoma; READ, rectum adenocarcinoma; SARC, sarcoma

These pan-cancer concept profiles formed shared blocks that reflected relationships among biologically related cancer types. For instance, low-grade glioma (LGG) and glioblastoma (GBM), both tumors of glial-cell origin^20^, exhibited highly similar concept profiles, differing primarily in cellularity, cellular atypia, and extent of necrosis (Fig. 3). Similarly, uterine corpus endometrial carcinoma (UCEC) and ovarian serous cystadenocarcinoma (OV) showed closely aligned profiles, consistent with their shared Müllerian epithelial origin and overlapping molecular features, particularly among serous subtypes involving hormonal and TP53-related pathways^21,22^. Colon adenocarcinoma (COAD) and rectal adenocarcinoma (READ), often jointly classified as colorectal adenocarcinoma of the lower gastrointestinal tract^23,24^, likewise displayed highly similar patterns characterized by gland-forming tumor epithelium with luminal necrosis. In contrast, kidney renal clear cell carcinoma (KIRC) and kidney renal papillary cell carcinoma (KIRP) exhibited distinct concept signatures, consistent with their divergent histogenesis from different nephron segments^25^ and distinct oncogenic driver programs associated with specific clinical behaviors and therapeutic implications^26,27^.

Closer inspection revealed both cancer-type-specific morphological signatures and shared pan-cancer concepts. Many concepts were selectively enriched within individual cancer types and corresponded to canonical diagnostic features. For example, breast invasive carcinoma (BRCA) was characterized by infiltrating tumor cells within fibrous stroma; GBM showed enrichment for densely cellular tissue with stroma; and kidney renal clear cell carcinoma (KIRC) exhibited tumor cells with characteristic clear, glycogen-rich cytoplasm. At the same time, several concepts recurred across multiple cancer types. Notably, patterns such as expanding thin fibrillary stroma and sheets of tumor cells with necrosis or tissue cracks were observed across cancers, suggesting shared morphological responses across diverse tumor contexts. In summary, these findings highlight that PICASSO captures the structured diversity of morphological features across cancers while preserving biologically meaningful relationships between cancer types.

### Auditing tumor detection model via concept control

We next used PICASSO to *audit* downstream clinical models by linking their predictions to interpretable morphological concepts (Fig 4a). Unlike traditional auditing approaches that provide “passive” inspection of model behavior, PICASSO enables active probing: it directly tests how specific morphological features drive predictions by modulating individual concept activations. This approach resolves key limitations of existing auditing tools. For instance, saliency maps highlight influential image regions, but these regions are often difficult to translate into semantically meaningful concepts^28,29^. Conversely, counterfactual image-based explanation methods can synthesize alternative images but lack the ability to isolate and manipulate individual morphological features in a controlled manner^17,30^. PICASSO instead operates at the level of semantic concepts, providing a precise and clinically meaningful framework for auditing pathology FMs.

**Fig. 4.**
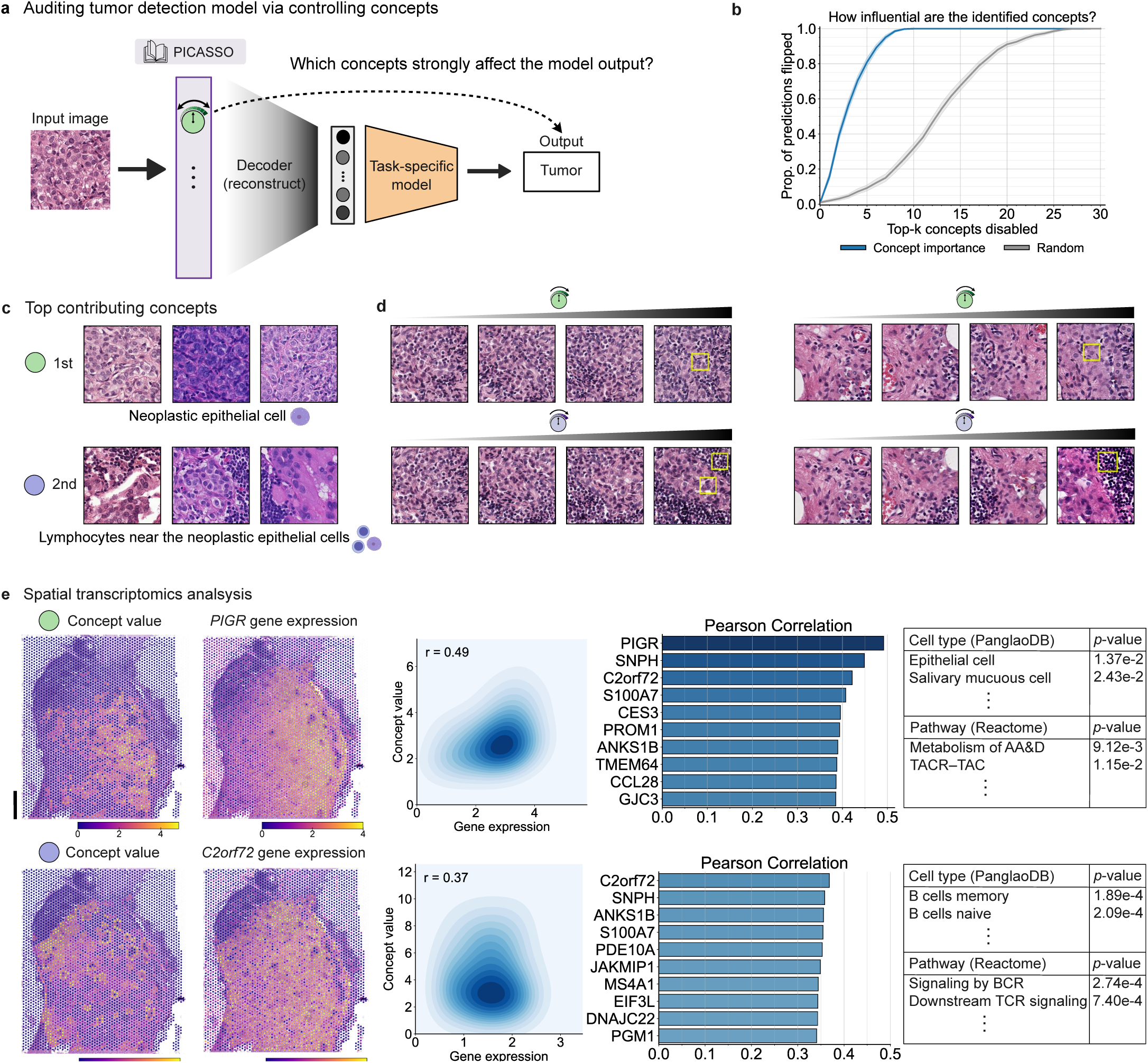
Identifying influential concepts for tumor detection model and studying their molecular signature. a, We use PICASSO to identify which concepts strongly affect the output of the tumor detection model, which gets FM embeddings as input. b, We benchmarked how accurately our method identified influential concepts. To evaluate, we conducted an ablation analysis on a set of 1,000 samples initially classified as a tumor. The y-axis plots the proportion of the samples whose prediction flipped to normal as the top-k most important concepts, ranked by Integrated Gradients, were progressively disabled. The random curve, where concepts were disabled in a random order, serves as a baseline. Lines represent the mean proportion across the samples, and shaded areas indicate the 95% confidence interval. c, Image patches that highly activate the top 1 and 2 most influential concepts are shown. d, For each top concept, we generate counterfactual images by progressively elevating the concept’s strength and visualizing the effect using Emb2Img. e, Spatial transcriptomics analysis highlights the biological relevance of the learned concepts. Left panels show the spatial distribution of concept activation values overlaid on tissue sections, which closely mirrors the expression patterns of top-correlated genes (*PIGR* for the 1st concept, *C2orf72* for the 2nd). Those genes show Pearson’s *r* = 0.49 and *r* = 0.37 with the concepts, respectively. Bar charts show the top correlated genes, and tables show the results of gene set enrichment analysis for all genes with Pearson’s *r >* 0.3. Black scale bar of WSI represents 1 mm.

As a case study, we audited a tumor detection model trained to identify metastatic breast cancer in lymph node tissue from the CAMELYON16 dataset^31^. To quantify concept importance, we applied Integrated Gradients^32^ to attribute model predictions to individual PICASSO concepts, rather than pixels or image regions, enabling attribution at a semantic, morphology-level resolution (Methods). To validate that these attribution scores capture the concepts most influential for tumor prediction, we performed a systematic ablation analysis: concepts were removed in descending order of importance, and the resulting impact on model output was measured. Strikingly, suppressing only the top five concepts flipped the prediction from “tumor” to “normal” in over 80% of tumor-classified images (Fig. 4b). These results indicate that model decisions rely on a sparse and identifiable set of morphological features captured by PICASSO.

We next examined the highest-ranking concepts consistently driving tumor predictions. Image patches strongly activating the top-ranked concept exhibited a canonical hallmark of metastatic carcinoma: cords and sheets of large neoplastic cells with enlarged nuclei and abundant cytoplasm, alongside scattered small lymphocytes in the background (Fig. 4c, top panel; Supplementary Fig. 7). To validate this interpretation, we progressively increased activation of this concept and visualized the resulting reconstructions using Emb2Img. This virtual intervention produced a marked increase in density of neoplastic cell content, confirming that the concept captures neoplastic cellularity (Fig. 4d, top panel).

The second-most influential concept captured a mixture of lymphocytes and neoplastic cells with enlarged and more hyperchromatic nuclei, both of which increased in parallel with concept activation (Fig. 4c-d, bottom panel; Supplementary Fig. 8), again recapitulating key morphological features of metastatic disease.

Together, these results show that the model identifies metastatic tumor cells in lymph nodes using clinically established morphological hallmarks, and that PICASSO reveals these decision cues in terms of interpretable pathological concepts. More broadly, this case study demonstrates how PICASSO can audit pathology AI systems and assess whether their predictions are grounded in clinically meaningful reasoning.

### Linking visual concepts to molecular biology

We next anchored PICASSO’s interpretable histopathological concepts in molecular biology by linking visual morphology to underlying gene-expression programs using spatial transcriptomics (ST) data from matched tissue sections, which provide paired histology images and gene expression measurements^33^ (Methods). This linkage enables biological interpretation, hypothesis generation, and the identification of candidate biomarkers and therapeutic targets. We applied this approach to the tumor-detection task using an ST dataset of metastatic breast cancer in lymph nodes, comprising 18,582 tissue spots from four patient samples^34^.

We first examined the molecular signature of the top-ranked tumor-detection concept, which visually corresponded to neoplastic cells (Fig. 4c, top panel). Correlating concept activation with gene expression across all ST spots revealed the strongest association with *PIGR1* (Pearson’s *r*=0.491), a gene involved in epithelial immunoglobulin transport and mucosal defense^35,36^. Spatial visualization on the whole-slide image (WSI) confirmed striking co-localization of concept activations and *PIGR* expression within the same tissue regions, consistent with its known role in epithelial immune transport^37^ (Fig. 4e and Supplementary Fig. 11).

To obtain broader biological insight, we performed gene set enrichment analysis (GSEA)^38^ on genes highly correlated with concept activation (Pearson’s *r >* 0.3) (Supplementary Table 2). Enriched terms included epithelial cell markers and pathways related to metabolism, Myc targets, and MAPK signaling, consistent with the neoplastic epithelial morphology captured by the concept. We complemented this analysis using frontier large language models (LLMs), which have recently been applied for gene set interpretation^39^ (Methods). The LLM consistently interpreted the correlated gene sets as reflecting epithelial tissue interacting with immune elements (Supplementary Fig. 9). Together, these results support that PICASSO’s neoplastic cell concept captures bona fide epithelial tumor signatures.

Similarly, the second-most influential concept, capturing a mixture of lymphocytes and neoplastic cells (Fig. 4c; *bottom*), was strongly correlated with the canonical B-cell marker *MS4A1* and candidate metastatic breast cancer markers such as *S100A7* ^40,41^ and *C2orf72* ^42^ (Fig. 4e, Supplementary Fig. 12, and Supplementary Table 3). GSEA further revealed enrichment for B cells and B-cell receptor signaling, while LLM-based interpretation additionally highlighted PI3K-AKT signaling (Supplementary Fig. 10), reflecting coordinated immune and tumor-associated programs.

Overall, integrating PICASSO with spatial transcriptomics reveals molecular programs underlying visually defined concepts, linking histomorphology to gene-expression structure. While demonstrated here in the well-characterized setting of metastasis detection, this framework extends to tasks where relevant morphology is less well understood, such as mutation prediction or survival modeling from H&E images, enabling discovery of biologically grounded morphological biomarkers.

### Identifying potential novel biomarkers associated with *EGFR* mutation

Determining *EGFR* mutation status in lung adenocarcinoma is critical for identifying patients who may benefit from tyrosine kinase inhibitor (TKI) therapy. Recently, pathology FMs have demonstrated the ability to infer *EGFR* mutations directly from routine H&E-stained images with high accuracy, including in prospective clinical studies^5,8^. This computational approach could complement standard clinical assays, such as polymerase chain reaction (PCR) and next-generation sequencing, by providing a rapid, cost-effective, and tissue-preserving alternative (Fig. 5a). However, the mechanisms underlying these predictions remain poorly understood: how do models detect molecular alterations that are not readily discernible to human pathologists?

**Fig. 5.**
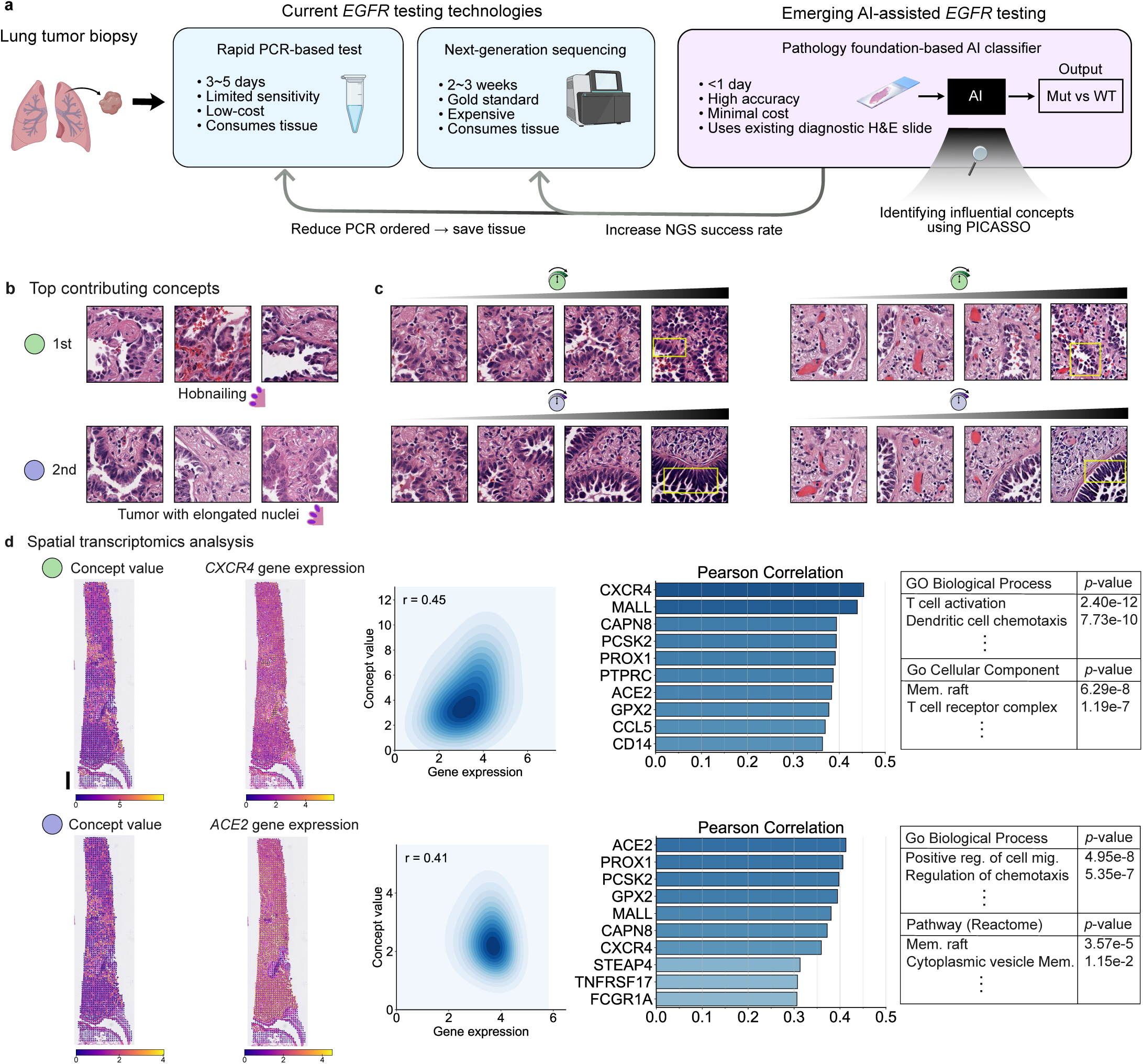
Identifying influential concepts for *EGFR* mutation status detection model and studying their molecular signature. a, Comparison of standard *EGFR* testing technologies with a rapid, cost-effective AI-assisted approach that infers the mutation status from routine H&E slides. We use PICASSO to render this “black box” model transparent. b, The top 1 and 2 most influential concepts for driving the model’s inference of *EGFR* mutation status. For each concept, image patches that highly activate the concept are shown. c, For each top concept, we generate counterfactual images by progressively elevating the concept’s strength and visualizing the effect using Emb2Img. d, Integration with spatial transcriptomics links these morphological concepts to molecular biology. Left panels show the spatial distribution of concept activation values overlaid on tissue sections, which closely mirrors the expression patterns of top-correlated genes (*CXCR4* for the 1st concept, *ACE2* for the 2nd). Bar charts show the top correlated genes, and tables show the results of gene set enrichment analysis for all genes with Pearson’s *r >* 0.2. Black scale bar of WSI represents 1 mm. Mut, Mutation; WT, Wildtype; Mem, Membrane; Reg, Regulation.

We used PICASSO to identify visual concepts driving FM-based prediction of *EGFR* mutation status. Two top-ranked concepts captured malignant epithelium and exhibited the classical “hobnailing” morphology^43^—epithelial tumor cells with apical protrusions bulging into glandular lumina (Fig. 5b). Visualization of concept activations across H&E WSIs revealed strong enrichment of both concepts in *EGFR*-mutant samples (Fig. 5b; Supplementary Fig. 13-14). Slide-level analysis across the TCGA LUAD cohort further confirmed significant associations between both concepts and *EGFR* mutation status (Pearson’s *r*=0.189, *p* = 1.3 × 10*^−^*^5^ and Pearson’s *r*=0.330, *p* = 6.7 × 10^−15^; *n* = 530 patients). Notably, the second concept, characterized by tapered and elongated point-like apical protrusions, was less prevalent but spatially more localized than the first.

We next perturbed each concept independently to characterize the morphological features driving these two concepts by progressively increasing concept activation and examining the resulting counterfactual morphologies (Fig. 5c). This analysis revealed subtle distinctions not apparent from exemplar patches alone. The first concept showed broad enrichment of tumor and stromal structures, encompassing hobnailing glands together with surrounding stroma (Fig. 5c, top panel). In contrast, the second preferentially modulated epithelial linings composed of elongated, hyperchromatic, and higher-grade malignant cells (Fig. 5c, bottom panel). Thus, although both concepts captured tumor microarchitecture, the second isolated more specific epithelial features. To our knowledge, these results provide the first direct visualization of histopathologic features underlying *EGFR* mutation prediction and reveal distinct morphological components associated with model inference.

We next investigated the molecular basis of these concepts by correlating concept activations with gene expression using lung adenocarcinoma spatial transcriptomics data (7,505 tissue spots from two patients)^33^ (Fig. 5d). The first concept was strongly associated with immune-related genes (Supplementary Table 4), including *CXCR4* (*r*=0.45), *MALL* (*r*=0.44), *CAPN8* (*r*=0.39), as well as *EGFR* itself (*r*=0.29), with enrichment for T-cell activation and antigen presentation pathways. The second concept showed strong associations with epithelial differentiation markers (Supplementary Table 5), including *ACE2* (*r*=0.41), *PROX1* (*r*=0.41), and *PCSK2* (*r*=0.40), together with enrichment for cell migration and membrane organization pathways. These findings suggest that the *EGFR* mutation-prediction model leverages cellular morphologies characterized by aberrant epithelial polarity and apical protrusion phenotypes, with the two concepts emphasizing slightly different aspects of tumor architecture.

Together, these results demonstrate that PICASSO, when integrated with matched spatial transcriptomic data, can link the morphologic features underlying FM-based mutation predictions to their associated gene-expression programs.

### Removing spurious artifact signals to mitigate shortcut learning

A major barrier to the safe clinical deployment of AI is shortcut learning, in which models exploit spurious, non-biological artifacts rather than true pathological features, thereby undermining reliability^44–47^. Because PICASSO learns concepts in an unsupervised manner, its concept atlas captures recurring visual patterns, including technical artifacts such as tissue folds introduced during sample preparation. This enables targeted suppression of artifact-associated concepts, reducing a classifier’s reliance on shortcuts while preserving biologically meaningful morphology. We identified five concept nodes corresponding to tissue-folding artifacts (Methods). Setting their activations to zero selectively removed the artifactual signals from FM embeddings. Visualization of this *concept intervention* using Emb2Img confirmed its effectiveness: reconstructed images showed elimination of tissue folds while preserving the underlying histopathology (Fig. 6a). Suppressing only the two dominant artifact concepts removed tissue folds in more than 80% of cases, as determined by manual inspection of reconstructed images (Fig. 6b).

**Fig. 6.**
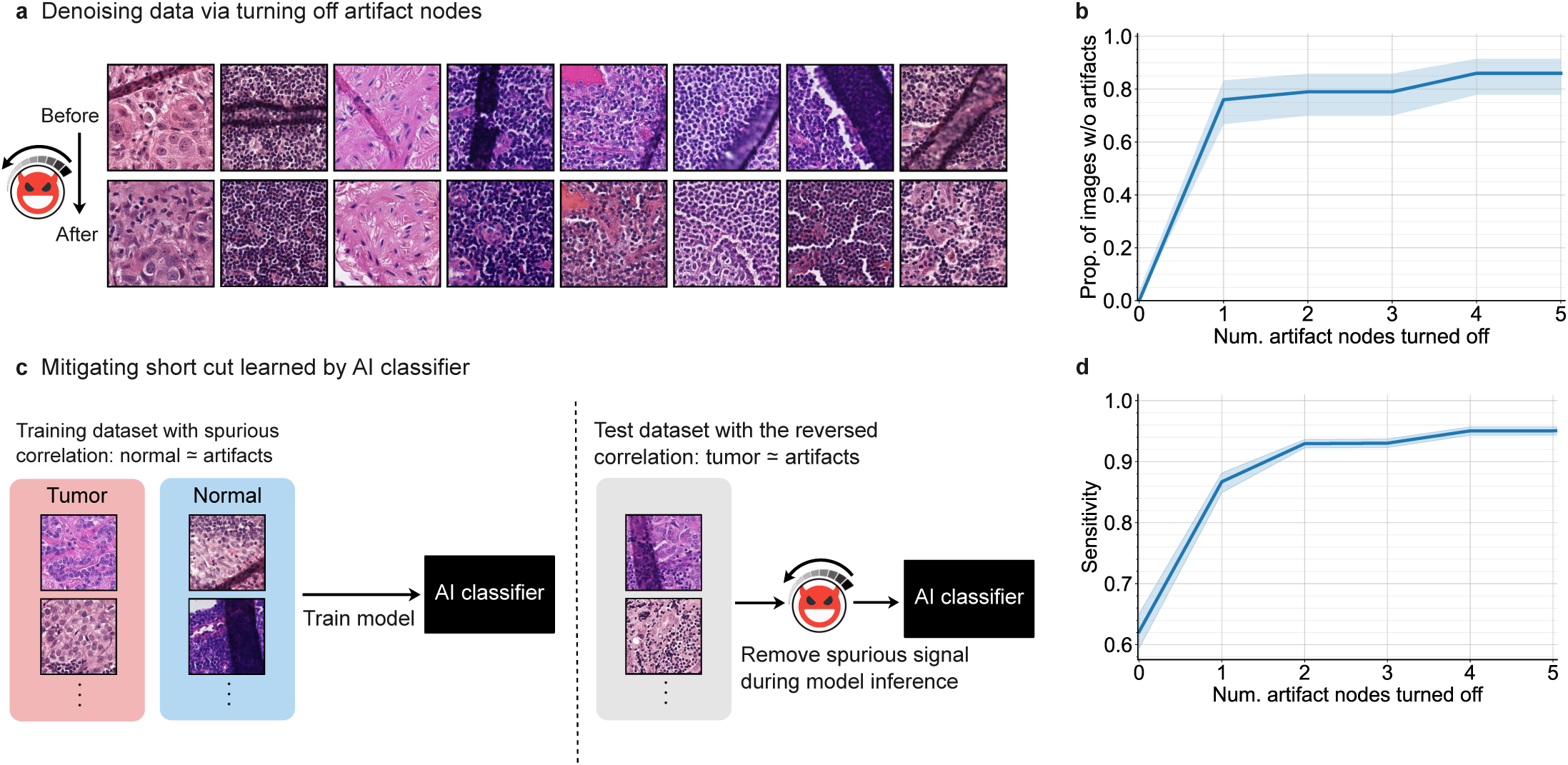
PICASSO corrects shortcut learning by selectively removing artifact concepts. a, Representative images showing the effect of PICASSO’s denoising procedure. Original images with tissue folding artifacts (top) are shown alongside their denoised counterparts (below), in which artifact-related concept nodes have been suppressed. b, Success rate of the denoising depending on the number of artifact nodes intervened on. The y-axis represents the proportion of images in which artifacts were successfully removed, evaluated on a sample of *n* = 100 images containing artifacts. The solid line denotes the observed proportion, and the shaded region represents the 95% Wilson score confidence interval. c, We simulate a shortcut learning setting in clinical AI development; a classifier is trained on a biased dataset where artifacts were spuriously correlated with the target label, but tested on a dataset with the opposite correlation. We can use PICASSO’s denosing capability to remove spurious signals during model inference. d, Sensitivity of the classifier on the external test set as a function of the number of artifact nodes intervened. Evaluations are repeated with *n* = 10 different train–test splits. The shaded areas indicate the 95% confidence interval, with the center line representing the mean.

We next tested whether this concept-based denoising strategy could correct a classifier’s reliance on a learned shortcut. To simulate this failure mode, we trained a tumor classifier on a biased dataset in which tissue folds were spuriously correlated with normal tissue and evaluated it on a test set in which the correlation was reversed, such that artifacts co-occurred with tumor tissue (Fig. 6c). As expected, the classifier learned the shortcut and failed to generalize: across *n* = 10 train-test splits, the classifier’s sensitivity—defined as the proportion of tumor samples correctly classified as tumor—remained high on the biased validation set (mean = 0.990) but dropped sharply on the external test set (mean = 0.612). In clinical settings, such degradation could translate into missed diagnoses. Notably, applying PICASSO’s denoising procedure at test time—without retraining—restored sensitivity to 0.950 (Fig. 6d).

These results demonstrate that PICASSO can expose and mitigate shortcut learning by selectively suppressing artifact-associated concepts while preserving diagnostically meaningful morphology, providing a principled path toward more interpretable, robust, and reliable clinical AI systems.

### Simulating therapeutic response via concept-guided interventions

PICASSO’s ability to manipulate clinically meaningful morphological features enables a new paradigm for clinical research: simulating therapeutic interventions through controlled modification of visual concepts, which we refer to as *virtual interventions*. By generating virtual tissue samples and evaluating outcomes using validated predictive models, this approach enables rapid *in silico* testing of therapeutic hypotheses, complementing the costly and time-intensive process of clinical trials.

As a proof of concept, we simulated the effects of tumor-infiltrating lymphocyte (TIL) therapy (Fig. 7a), an FDA-approved immunotherapy^48^ in which a patient’s T cells are expanded *ex vivo* and reinfused to enhance anti-tumor immunity. We identified ten PICASSO concepts corresponding to lymphocytes (see Methods) and digitally increased TIL abundance within tumor regions of TCGA WSIs from breast invasive carcinoma patients by progressively amplifying these concept activations (Fig. 7b). Visual inspection confirmed increased lymphocyte density within tumor regions while preserving the underlying neoplastic architecture.

**Fig. 7.**
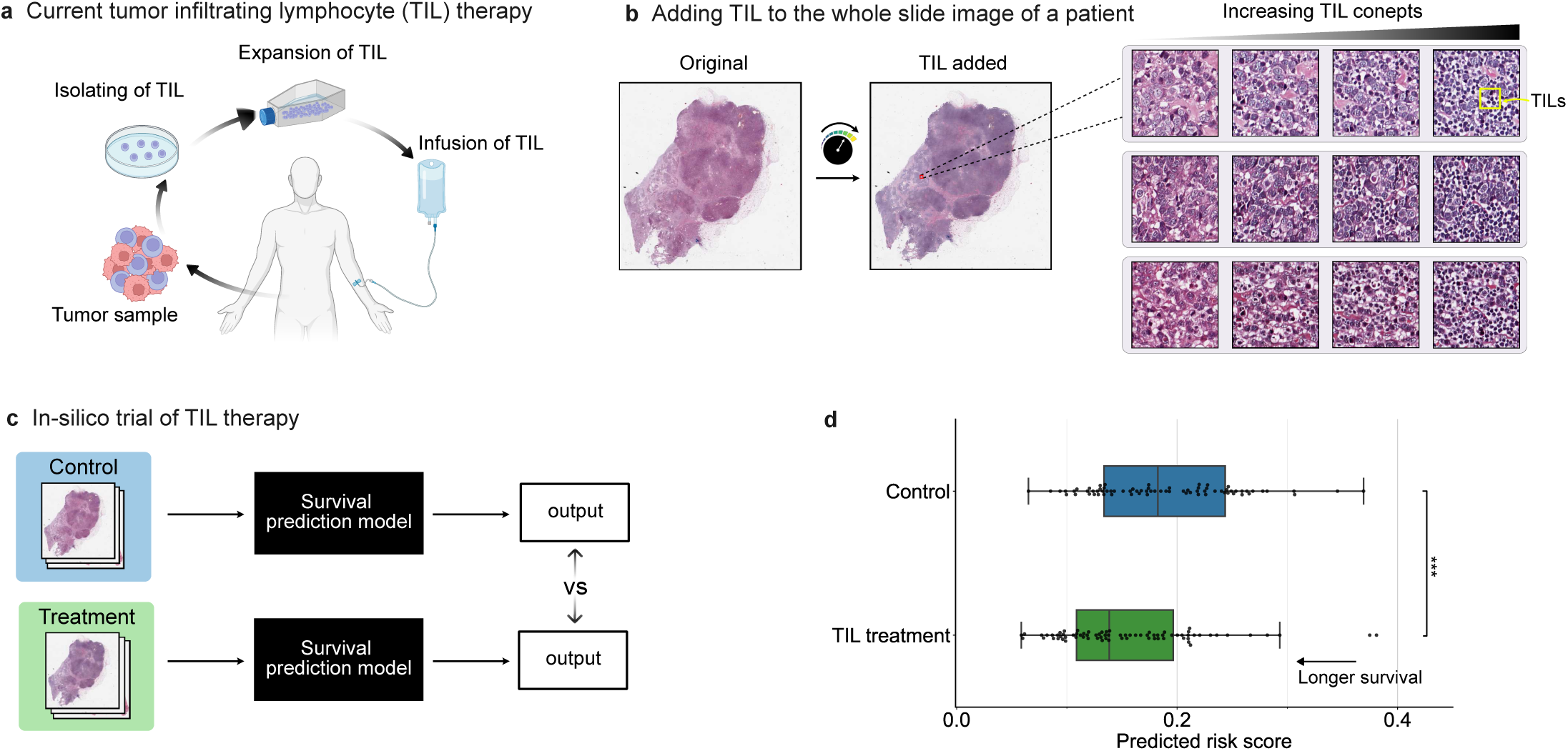
Simulating TIL therapy using PICASSO. a, Current scheme of tumor-infiltrating lymphocyte therapy. b, We use PICASSO to simulate the morphological effects of TIL therapy. After we increase the TIL concepts, we visualize the modulation effects by visualizing the reconstructed embeddings using Emb2Img. As we increase the TIL concepts, more lymphocytes appear in the image. c, Prognosis is predicted using survival models, and predictions are compared between the control (original) and the treatment (TIL enriched). d, Comparison of predicted survival risk scores between the control (original patient images) and TIL treatment (computationally TIL-enriched) groups. The boxes represent the interquartile range (IQR) with their lower and upper bounds corresponding to the first and third quartiles, respectively, and the center lines indicate the medians. The whiskers extend to 1.5×IQR from the box. Each dot represents the predicted risk score for an individual patient. We performed a two-sided Mann-Whitney U test to assess the statistical significance of the difference between the two distributions. (***: *p <* 0.001)

We next evaluated the predicted effects of this virtual intervention using a survival prediction model trained with the PORPOISE framework^49^, which achieved a concordance index of 0.705 ± 0.063, consistent with the original study (Supplementary Figs. 1-2; Methods). Comparing model outputs between the original WSIs (control) and TIL-enriched counterparts (treatment) in the test cohort (*n* = 76), we observed a significant reduction in predicted risk scores following TIL augmentation (Mann-Whitney *U* = 3, 827, p-value = 4.35 × 10^−31^) (Fig. 7c).

These results demonstrate that PICASSO enables targeted and clinically meaningful virtual interventions on digital pathology images, providing a platform for conducting *in silico* clinical trials. While not a replacement for prospective studies, this framework provides a scalable preclinical approach for exploring therapeutic hypotheses, probing morphology-outcome relationships, and prioritizing patient subgroups or treatment strategies for further investigation.

## Discussion

### Motivation

Pathology foundation models (FMs) have substantially advanced computational pathology in recent years, achieving major performance gains across a broad range of clinical tasks^4–6,8,18^. Despite these advances, a fundamental challenge remains: pathology AI systems built on FM embeddings remain largely opaque, limiting interpretability, biological insights, and confidence in clinical deployment.

### PICASSO overview

Here, we present PICASSO, a framework that makes pathology FM embeddings interpretable and controllable by reorganizing learned representations into a concept space of human-interpretable morphological features. PICASSO constructs, to our knowledge, the first pan-cancer atlas of histomorphological concepts derived from more than 120 million image patches spanning 32 cancer types. Across diverse downstream applications, PICASSO enables auditing of model behavior, biological discovery, mitigation of shortcut learning, and simulation of therapeutic interventions.

A key innovation of PICASSO is its integration with Emb2Img model, a diffusion-based model that maps FM embeddings back to realistic histopathological images. Unlike traditional “passive” explainability approaches that rely on static exemplars, PICASSO enables dynamic visualization through controlled modulation of individual concepts and observation of the resulting morphological changes. This capability links abstract embedding representations to clinically meaningful histopathologic phenomena and addresses longstanding limitations of prior dictionary-learning^10^ and feature-attribution methods^28,29,50^.

### EGFR auditing

PICASSO reframes explainable AI in pathology from a retrospective auditing tool into a virtual platform for scientific discovery. In an *EGFR* mutation-prediction model, PICASSO revealed that predictions were consistently driven by a subtle “hobnailing” epithelial morphology—a feature not previously recognized as associated with *EGFR* status by pathologists. Integration with spatial transcriptomics further linked this morphology to epithelial signaling gene expression programs, illustrating how PICASSO can refine morphologic taxonomies, generating biologically grounded hypotheses, and identify candidate biomarkers that merit independent clinical validation.

### Artifact removal

Beyond interpretation, concept-level manipulation provides a principled solution to shortcut learning, in which models exploit technical artifacts rather than true pathology. PICASSO enables selective suppression of artifact-associated concepts during inference without retraining. By identifying and suppressing tissue-fold concepts, we restored the sensitivity of metastasis classifiers from 0.61 to 0.95 in an external cohort in which the spurious correlation was reversed, demonstrating that targeted concept manipulation can both diagnose and mitigate model vulnerabilities.

### In-silico trial

PICASSO further enables controlled counterfactual experimentation that is difficult to achieve in conventional pathology studies. Observational cohorts are constrained by sample size and confounding, but by modulating clinically relevant concepts–such as lymphocyte infiltration—in representation space, we simulated therapeutic interventions and estimated downstream effects on patient outcomes. Although dependent on the fidelity of downstream survival models and not a substitute for prospective clinical trials, this framework provides a scalable approach for testing mechanistic hypotheses, prioritizing interventions, and refining trial design in settings where direct experimentation is impractical.

### Outlook

Beyond the scope of this study, several extensions and limitations motivate future work. Although we focused on histopathology, a central modality in medicine, the principles underlying PICASSO may extend to FMs in other biomedical modalities, including radiology, genomics, and retinography, where model capabilities have advanced rapidly but interpretability remains limited^51,52^. In addition, although we linked histologic concepts to transcriptomic programs, emerging multimodal datasets, such as high-plex proteomics^53–55^, may further refine concept definitions and their biological grounding. The morphologic biomarkers identified here, such as the hobnailing feature associated with *EGFR* prediction, will also require validation in independent clinical cohorts before translation to patient care. Likewise, *in-silico* clinical trials depend on how accurately downstreamprediction models capture complex biological relationships and therefore cannot replace prospective clinical studies. Finally, concepts in PICASSO vary in abstraction level, and extending the framework toward hierarchical concept representations may better capture the multi-scale organization of tissue morphology.

### Impact summary

PICASSO provides actionable capabilities across the pathology AI ecosystem, enabling developers to diagnose and correct model failures through concept-level perturbations, clinicians to gain transparency into model reasoning, biologists to formulate hypotheses grounded in molecular programs, therapeutic researchers to connect visual phenotypes with candidate drug targets, and regulators to systematically audit model behavior. By transforming opaque FM embeddings into interpretable and controllable representations, PICASSO establishes a foundation for more transparent, biologically grounded, and clinically reliable pathology AI. More broadly, this work illustrates how foundation models, when coupled with concept-based interpretability frameworks such as PICASSO, can evolve from black-box predictive systems into scientific instruments for probing disease mechanisms and advancing precision oncology.

## Methods

### PICASSO concept atlas learning

#### Data curation and preprocessing

We derived the pan-cancer dataset from 11,603 hematoxylin and eosin (H&E)-stained whole-slide images (WSIs) across 32 cancer types from The Cancer Genome Atlas (TCGA), accessed via the Genomic Data Commons portal (Supplementary Table 1). We preprocessed these WSIs by segmenting tissue regions from the background using the OpenSlide library^56^ and the CLAM toolkit^1^. From these tissue masks, we then extracted 153,889,118 non-overlapping, which serve as the basic processing unit for pathology AI models. Each patch measured 256 × 256 pixels at 20×-equivalent magnification (0.5 µm/px), corresponding to a physical size of 128 × 128 µm per patch. To prepare for concept atlas learning, we then split the resulting patch dataset into training (80%), validation (10%), and testing (10%) sets, ensuring that patches from the same slide were confined to a single split to prevent data leakage.

#### Sparse autoencoder for concept decomposition

Pathology foundation models (FMs) represent histopathology image patches as dense embeddings that are not directly interpretable. To decompose these embeddings into human-interpretable morphological concepts, we developed PICASSO, a framework based on a sparse autoencoder (SAE), which functions as a form of dictionary learning. The SAE consists of two main components:

**Encoder** Given an FM embedding *z_i_* ∈ R*^d^*, the encoder projects it into a much higher-dimensional latent space. Crucially, sparsity is enforced by retaining only the top-*k* activations:

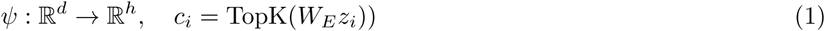

where *W_E_*∈ R*^D×d^* is the encoder matrix, and *c_i_* is the sparse concept representation satisfying a predefined sparsity level of ∥ *c_i_* ∥ _0_ = *k*.

**Decoder** The decoder reconstructs the original embedding (*z*^*_i_*) from the sparse representation:

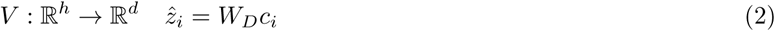

where *W_D_* ∈ R*^d×h^* is the decoding matrix–Bias terms are included in the encoder and decoder projections but omitted from the notation for clarity. Although the latent dimension *h* is set much larger than the original embedding dimension *d*, the reconstruction task remains non-trivial because sparsity is enforced at the encoding stage. Each non-zero entry in the sparse vector *c_i_* corresponds to an activated “concept node,” which encodes interpretable morphological concepts. Our implementation is based on SAELens, the open-source sparse autoencoder codebase^57^.

We applied this framework to the TCGA pan-cancer patch dataset using embeddings generated by Virchow2^4,18^, a state-of-the-art pathology FM that produces *d* = 1,280-dimensional embeddings. To ensure a rich concept vocabulary, we chose an expansion ratio of 32, yielding a latent dimension of *h* = 40,960. We determined the optimal level of sparsity by training the SAE across multiple sparsity levels (*k* ∈ {10, 30, 50, 80, 100}). We ultimately selected *k* = 30, as this provided the highest sparsity while maintaining strong reconstruction performance (explained variance >0.8). This configuration was used for all subsequent concept-driven analyses.

#### Optimization

We optimized SAE by minimizing the mean squared error (MSE) between the original and reconstructed embeddings: 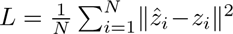 where 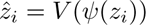. This objective ensured that the sparse sets of concepts faithfully represent information captured by the original FM embedding. We trained SAE for five epochs using the Adam optimizer^58^ with a learning rate of 1e-4 and a batch size of 2,048. Training one SAE took 14 hours and 10 minutes on a single NVIDIA A40 graphics processing unit (GPU).

### Visualzing FM embeddings with Emb2Img

To visualize how PICASSO concepts influence the pathology FM embeddings, we used Emb2Img, a conditional diffusion framework designed to map FM embeddings back into the image space. This allowed us to visualize the morphological effects of embedding perturbations directly—for example, when modulating a single concept node—a technique critical for concept verification.

The architecture of Emb2Img follows that of Stable Diffusion v2^59^, a text-to-image generative model, but with one key modification: image generation is conditioned on FM embeddings rather than text. For computational efficiency, the diffusion process operates in the latent space of a pretrained variational autoencoder (VAE)^60^, which compresses 512 × 512 images into 64 × 64 latent representations. All pathology images were first resized to match this VAE input resolution. The image generation process starts from Gaussian noise, and a U-Net^61^ denoiser progressively reconstructs clean VAE latents. Conditioning on FM embeddings is achieved via a cross-attention mechanism between the embeddings and the U-Net features. During generation, we applied classifier-free guidance (scale = 6.5) to ensure that the generated image faithfully reflects the conditioning embeddings. We used the PNDM scheduler for the diffusion process, with 50 denoising steps during training and 100 steps during inference.

Emb2Img was trained on the same extensive collection of patch–embedding pairs used to build the PICASSO concept atlas. Training followed the standard diffusion objective: progressively adding noise to clean VAE latents and optimizing the U-Net to predict this noise using MSE loss. We used the AdamW optimizer^62^ (learning rate = 1e-4) with gradient clipping (max norm = 1.0) and dropout on guidance conditioning (probability = 0.1). An exponential moving average (EMA) of model weights was maintained to reduce noise in the training process. The model was trained for 16,000 optimizer steps with an effective batch size of 1,024 (per-GPU batch size = 16, gradient accumulation = 16, across 4 NVIDIA A40 GPUs). Mixed-precision (fp16) training was used to improve memory efficiency and throughput. We employed PyTorch Distributed Data Parallel (DDP) for multi-GPU training. The total training time was 5 days and 22 hours.

#### Concept interpretation

To ensure the concept nodes learned by PICASSO were clinically meaningful and human-recognizable, we established a systematic interpretation and validation protocol in close collaboration with two board-certified pathologists. This process was essential for grounding the PICASSO concepts in established histopathologic terminology.

First, for initial annotation, we presented the image patches that maximally activated each concept node across our pan-cancer test set. The expert pathologists were tasked with examining these exemplars to identify a common, dominant morphological pattern and describe it using standard histopathological terminology. This analysis allowed us to assign preliminary labels to concepts representing clear features such as “dense lymphocytes,” “glandular structures,” or “‘spindle cell arrangements.”

Second, to refine the interpretations of more subtle concepts, we employed dynamic visualization using our Emb2Img model. For a given concept, we systematically increased its activation value for an image while holding others constant and generated a sequence of images where the concept was progressively exaggerated. Pathologists reviewed these sequences to observe the specific morphological transformation controlled by the concept. This interactive method was essential for defining the precise semantic axis of each concept and provided robust validation of its direct link to a specific histopathological feature.

### Pan-cancer concept profiling

We leveraged the capability of the PICASSO framework to extract and quantify morphological concepts from pathology image embeddings, enabling us to generate a comprehensive concept profile of TCGA samples across cancer types. We profiled concept activations across 6,812 whole slide images (WSIs) from 14 cancer types in TCGA, including 10 largest cancer types by sample size and 4 additional types chosen for their biological and morphological relevance. For each WSI, we first performed tumor region localization to focus our analysis on the region of interest. To achieve that, we used the CONCH vision-language model^63^, which assigns a cancer presence score to each patch based on the cosine similarity between patch embeddings and predefined tumor-related text prompts (Supplementary Table 6-7). To ensure a focus on highly probable tumor regions, we restricted our analysis to the top 10% of cancer-scored patches for each WSI. Within this tumor area, we then computed the mean activation of all PICASSO concepts. For generating the visualization-friendly heatmap, we selected concepts as follows. We first retained the top 1,000 concepts based on their global mean activation across all cancer samples. Among them, we highlighted the top five most active concepts specific to each individual cancer type.

### Training classifier

We trained and evaluated downstream models on a range of clinical tasks using WSIs from distinct sources: TCGA, CAMELYON16, and EBRAINS. For all tasks, we first precomputed embedding of patches extracted at the x20-equivalent magnification level for three leading pathology FMs: Virchow2 (d=1,536), UNI2 (d=1,536)^6^, GigaPath (d=1,536)^5^. In addition, for comparison, we computed embeddings of a ResNet50 model (d=2,048)^64^, extracted from the penultimate layer of the model, which was pre-trained on ImageNet.

#### Attention-based multiple instance learning

Given that most pathology prediction tasks provide only slide-level labels, we employed attention-based multiple instance learning (ABMIL) models^29^. This architecture is essential as it enables weakly supervised training using only slide-level labels, eliminating the need for costly patch-level annotations. The ABMIL approach simultaneously learns to: (1) assign attention weights to individual patches, thereby prioritizing the most relevant regions for the downstream task, and (2) predict the label using a slide-level representation constructed via a weighted average of patch embeddings. Specifically, for a slide *i* with *n_i_* patch embeddings 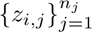, the preicted label 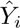 is computed as follows:

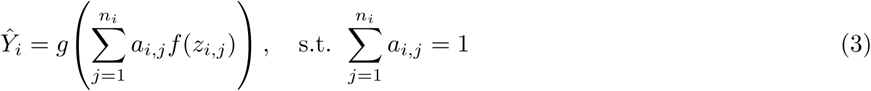

where *a_i,j_* denotes attention weights assigned by the gated attention mechanism to the embedding *z_i,j_*, *f* (·) is the patch embedding transformation function, and *g*(·) is the final prediction head. For classification tasks, models were trained using the cross-entropy loss. For prognosis prediction tasks, models were trained using the specialized time-to-event modeling approach detailed in the following section. All models were optimized using Adam^58^ (learning rate 1e-4) with a batch size of one WSI.

#### Survival modeling

To perform prognosis prediction, we had to adapt the ABMIL architecture for survival prediction; We made changes to the data processing and loss function to accommodate time-to-event and censoring data. Specifically, we used the discrete-time survival modeling approach^49^, which frames the problem as a multi-task ordinal regression. Patient survival time was discretized into one of four time bins, defined by the quartiles of event times in the training cohort: [0, q1), [q1, q2), [q2, q3), and [q3, ∞). The ABMIL model was configured to output a four-dimensional vector, representing the discrete-time hazard probabilities across these intervals. Each predicted hazard represents the conditional probability of an event occurring in interval *j*, given survival up to that interval. The final survival function *S*(*t*), which is the probability of surviving beyond time *t*, is then calculated as the cumulative product of the survival probabilities across all preceding time intervals. The model was trained using a negative log-likelihood loss tailored for right-censored survival data. We applied this framework to train separate survival models for breast invasive ductal carcinoma (TCGA–BRCA) and skin cutaneous melanoma (TCGA–SKCM).

### Downstream clinical tasks for training task-specific models

#### Breast metastasis detection on CAMLEYON16

For breast cancer metastasis detection, we used the Cancer Metastases in Lymph Nodes Challenge 2016 (CAME-LYON16) dataset^65^, which contains 400 whole-slide images (WSIs) of sentinel lymph nodes collected at Radboud University Medical Center and University Medical Center Utrecht. Each slide was annotated for tumor presence by expert pathologists. We applied 10-fold Monte Carlo cross-validation with partitioning at the patient level. The official test set was retained, and the remaining slides were divided into training and validation sets in an 80:20 ratio. For each fold, the model with the lowest validation loss was selected for evaluation.

#### Non-small cell lung cancer subtyping on TCGA

For non-small cell lung cancer (NSCLC) subtyping, we used lung adenocarcinoma (LUAD) and lung squamous cell carcinoma (LUSC) samples from TCGA. Data were split at the patient level into training, validation, and test sets in an 80:10:10 ratio. We performed 10-fold Monte Carlo cross-validation, selecting the best model in each fold based on the area under the receiver operating characteristic curve (AUROC).

#### BRCA receptor status prediction on TCGA

For receptor status prediction (HER2, PR, ER), we used breast cancer (BRCA) slides from TCGA. Because labels are defined at the patient level, a single representative slide was retained for each patient: if multiple diagnostic slides (DX1) were available, we selected either the DX1 slide with the largest tissue area or, if only one DX1 slide was present, that slide. Data were partitioned at the patient level into training, validation, and test sets in an 80:10:10 ratio, with 10-fold Monte Carlo cross-validation. The best-performing model in each fold was chosen using AUROC.

#### Microsatellite instability prediction

For microsatellite instability (MSI) prediction, we used colon and rectal adenocarcinoma (COAD/READ) WSIs from TCGA. When multiple slides were available per patient, only one representative DX1 slide, or the DX1 slide with the largest tissue area, was retained. Patient-level partitioning was used to split the data into training, validation, and test sets in an 80:10:10 ratio. Model development followed 10-fold Monte Carlo cross-validation, and AUROC was used for model selection.

#### *EGFR* mutation prediction

For epidermal growth factor receptor (EGFR) mutation prediction, we used lung adenocarcinoma (LUAD) WSIs from TCGA. Only one diagnostic slide (DX1) per patient was retained, or the DX1 slide with the largest tissue area if multiple were available. To account for the low prevalence of the positive class, data were partitioned at the patient level into training, validation, and test sets in a 60:20:20 ratio. We used 10-fold Monte Carlo cross-validation, with AUROC as the model selection criterion.

#### IDH mutation on EBRAINS

For IDH mutation prediction, we used brain tumor WSIs from the EBRAINS dataset, comprising 795 patients and 873 specimens with annotated IDH status (540 wildtype and 333 mutant). The dataset included 508 glioblastomas, 189 astrocytomas, and 176 oligodendrogliomas. Data were split at the patient level into training, validation, and test sets in an 80:10:10 ratio. We used AUROC for model selection across 10-fold Monte Carlo cross-validation.

#### Prognosis prediction

For prognosis modeling, we used breast invasive carcinoma (BRCA) and cutaneous melanoma (SKCM) WSIs from TCGA. Overall survival times and censoring information were obtained from the Genomic Data Commons. For patients with multiple slides, we retained one diagnostic DX1 slide or, if multiple DX1 slides were available, the slide with the largest tissue area. Data were split at the patient level into training, validation, and test sets in an 80:10:10 ratio. Models were trained with 10-fold Monte Carlo cross-validation, and the concordance index was used for model selection.

### Auditing classifier via PICASSO concepts

To provide a faithful, concept-level explanation for the predictions made by the downstream classifiers, we employed the Integrated Gradients (IG) method^32^, an axiomatic attribution technique. IG was originally developed for attributing a classifier’s output to input-level features such as pixels. When adapted to concept-level features, this technique allows us to quantify the contribution of each PICASSO concept node to the final prediction, effectively attributing the classifier’s decision back to specific morphological features. Specifically, we attribute the prediction, represented as *G*(*c_i_*) = *g*(*ψ*(*c_i_*)), back to the sparse concept representation *c_i_*. The IG value for the *j*-th concept node is calculated as:

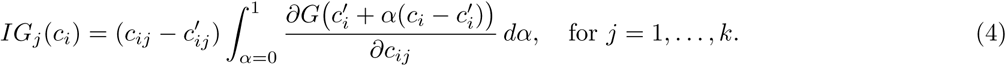

where *c^′^* represents the baseline concept vector, which we set to the null vector (i.e., *c^′^* = 0). By the completeness axiom, IG ensures that the attributions sum to the prediction difference, i.e.,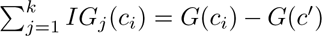, meaning that the total “credit” assigned to concepts fully accounts for the model’s output. To identify concepts with global importance across the entire dataset, we computed the average Integrated Gradient, *IG_j_*, for each concept node *j* across all analyzed samples.

### Processing spatial transcriptomics

To investigate the relationship between the histopathological concepts identified by PICASSO and the underlying molecular biology, we integrated our analysis with spatial transcriptomics (ST) data. We obtained preprocessed ST profiles from the Hest-1K dataset^33^, a comprehensive collection of 1,229 spatial transcriptomic profiles spanning 25 cancer types. An advantage of this dataset is that it has already been preprocessed to ensure that ST spots are spatially aligned with their corresponding WSIs precisely, providing a link between tissue morphology and transcriptomics measurements. We collected ST data samples for tissue and cancer types matching that of the downstream task dataset of interest. For the metastasis detection model, we used 10x Genomics Visium profiles of four metastatic lymph node samples from breast cancer patients^34^. For the *EGFR* detection model, we used 10x Genomics Xenium profiles from two patients with lung adenocarcinoma.

Prior to analysis, we performed quality control on the ST data. We filtered out genes that were expressed in less than 10% of the spatial spots. Subsequently, we normalized the counts per cell and applied a log(1+*x*) transformation. Following this preprocessing, we identified highly variable genes using the ‘highly variable genes’ function implemented in the scanpy package^66^. Our analysis focused on this identified set of highly variable genes. To identify specific genes associated with a particular morphology represented by a PICASSO concept, we computed the Pearson correlation coefficient between the activation of each concept node and the expression level of each highly variable gene across all ST spots.

For each concept, we selected the top correlated genes and performed gene set enrichment analysis with Enrichr^38^, which enables enrichment testing against a broad collection of curated gene set databases. In addition to database-driven enrichment, we used a large language model (LLM) to generate complementary, free-text functional hypotheses from the concept-associated gene lists. Prior work has shown that LLMs can summarize gene set function and provide mechanistic rationales, particularly when curated databases provide limited coverage^39^. Specifically, we used OpenAI GPT-5, building on prior evidence that GPT-4–class models perform well for gene set interpretation. We prompted the LLM with the concept-associated gene list and requested it to provide a general overview, functional clusters, pathway-level insights, and summary interpretation. We treated LLM-derived outputs as hypothesis-generating and reported them alongside the Enrichr results.

### Artifact removal

We leveraged the capability of the PICASSO framework for controlling morphological concepts to perform artifact removal, specifically targeting tissue folds, which are prevalent artifacts in histopathology. Our first step was to identify concept nodes that represent these artifacts. To this end, we trained a logistic regression with *L*_1_ penalty on the PICASSO concept activations to classify patches as either “artifact” vs “clean”. The model was trained using 2,000 manually labeled samples (1000 artifacts and 1000 clean). We explored the regularization strength hyperparameter *C* ∈ {0.003, 0.004, 0.005, 0.01, 0.02, 0.05}. From models that achieved a validation AUROC over 0.9, we closely examined the concept nodes that received non-zero weights. To verify their representation of tissue folds, we utilized the Emb2Img model by modulating the activation of each candidate concept node and observing whether this perturbation affected the presence or prominence of a tissue fold in the generated image. Through this process, we identified five key nodes that strongly represented tissue fold artifacts. Finally, we utilized these five nodes for the task of artifact removal. For any patch embedding to be denoised, we first decomposed it into its PICASSO concept activations. Any activation corresponding to one of the five identified artifact nodes was then selectively suppressed (set to zero). The resulting “denoised” embedding was subsequently reconstructed using the SAE decoder. The effectiveness of this concept-based artifact removal process was further visualized by applying the Emb2Img model to the denoised embeddings.

### In-silico clinical trial

To simulate TIL therapy by modulating the amount of lymphocytes, we first identified PICASSO concept nodes that represent lymphocytes. To that end, we trained a logistic regression model with *L*_1_ regularization on 100,000 patches (97,754 positive v.s. 2,246 negatives) that were independently annotated by a lymphocyte detector^65^. From the models that achieved a validation AUROC over 0.9, we selected the model exhibiting the highest sparsity. Concepts with nonzero weights were validated through perturbing the nodes and visualizing their effect using the Emb2Img model. Five nodes most strongly associated with lymphocytes were selected. We then simulated lymphocyte infiltration by increasing the strengths of the five nodes up to 10 within tumor regions identified by CONCH^63^. We then compared the predicted risk score determined by the survival prediction model, between the original slides (control) and the TIL-enriched slides (treatment). Survival curves were reconstructed from predicted hazard probabilities via cumulative survival across discrete time bins.

### Statistical analysis

For downstream clinical classification tasks, the patient served as the primary unit of analysis, with one representative diagnostic whole-slide image retained per patient. Model performance was evaluated using 10-fold Monte Carlo cross-validation; results are reported as means with 95% confidence intervals. Classification accuracy was assessed using the AUROC, survival prediction using the C-index, and Kaplan–Meier survival curves were compared using the two-sided log-rank test. For the in-silico clinical trial, differences in predicted risk scores between control and TIL-enriched groups were assessed using the two-sided Mann–Whitney U test. Associations between concept activations and gene expression (analyzed at the spot level) or slide-level mutation status were assessed using Pearson’s correlation coefficient. To account for multiple testing, *p* values were adjusted using the Benjamini–Hochberg procedure. For artifact suppression experiments, sensitivity was averaged across 10 independent train–test splits; Confidence intervals (CI) for binomial proportions (e.g., artifact removal success) were calculated using the Wilson score interval.

## Data availability

We obtained the data for building PICASSO from the TCGA Research Network. Raw whole-slide images (WSIs) and associated clinical metadata are available from the NCI Genomic Data Commons (https://portal.gdc.cancer.gov). Evaluation datasets used in this study are publicly accessible: CAMELYON16 (https://camelyon16.grand-challenge.org/); EBRAINS neuropathology collection (https://doi.org/10.25493/WQ48-ZGX); and HEST-1K spatial transcriptomics (https://huggingface.co/datasets/MahmoodLab/hest).

## Code availability

The code to run the PICASSO framework will be released for academic purposes once the article is published. Third-party code and pretrained models used in this work are publicly available at the following locations: SAELens sparse autoencoder codebase (https://github.com/jbloomAus/SAELens); UNI2 (https://huggingface.co/MahmoodLab/UNI2-h); GigaPath (https://huggingface.co/prov-gigapath/prov-gigapath); Virchow2 (https://huggingface.co/paige-ai/Virchow2); and the lymphocyte detector model (https://huggingface.co/kaczmarj/pancancer-lymphocytes-inceptionv4.tcga). We have documented all technical deep learning methods and software libraries used in the study while ensuring the paper is accessible to the broader scientific audience.

## Acknowledgements

C.K. and S.-I.L. received support from the National Institutes of Health (grants R01 AG061132, R01 EB035934 and RF1 AG088824). P.K.K. received support from the Simons Center for Quantitative Biology at Cold Spring Harbor Laboratory. Z.Z was supported in part by Developmental Funds from the Cancer Center Support Grant P30CA045508.

## Author contributions

C.K., P.K., and S.-I.L. conceived the study. C.K. and S.-I.L. developed the central concept of extracting human-interpretable concepts from pathology images and performing concept-based image editing, and established the overall methodological framework. C.K. led the implementation of the framework, designed and performed the experiments, conducted the analyses, implemented the codebase, and generated the figures. S.-I.L. identified cancer pathology as the primary application area and proposed integrating image concepts with transcriptomic analyses. J.K. contributed to pathology dataset acquisition, preprocessing, and domain-specific guidance. P.K. and S.-I.L. provided guidance on pathology AI applications, downstream analyses, experimental design, and presentation of results. Z.Z. and D.S. provided clinical pathology expertise, annotated and interpreted pathological concepts, analyzed pathology images, and helped validate the biological findings. C.K., Z.Z., P.K., and S.-I.L. interpreted the results and wrote the manuscript. S.-I.L. secured funding. P.K. and S.-I.L. provided senior supervision of the study. All authors reviewed and approved the final manuscript.

