## Supplementary Information for "Dissecting and directing pathology foundation models"

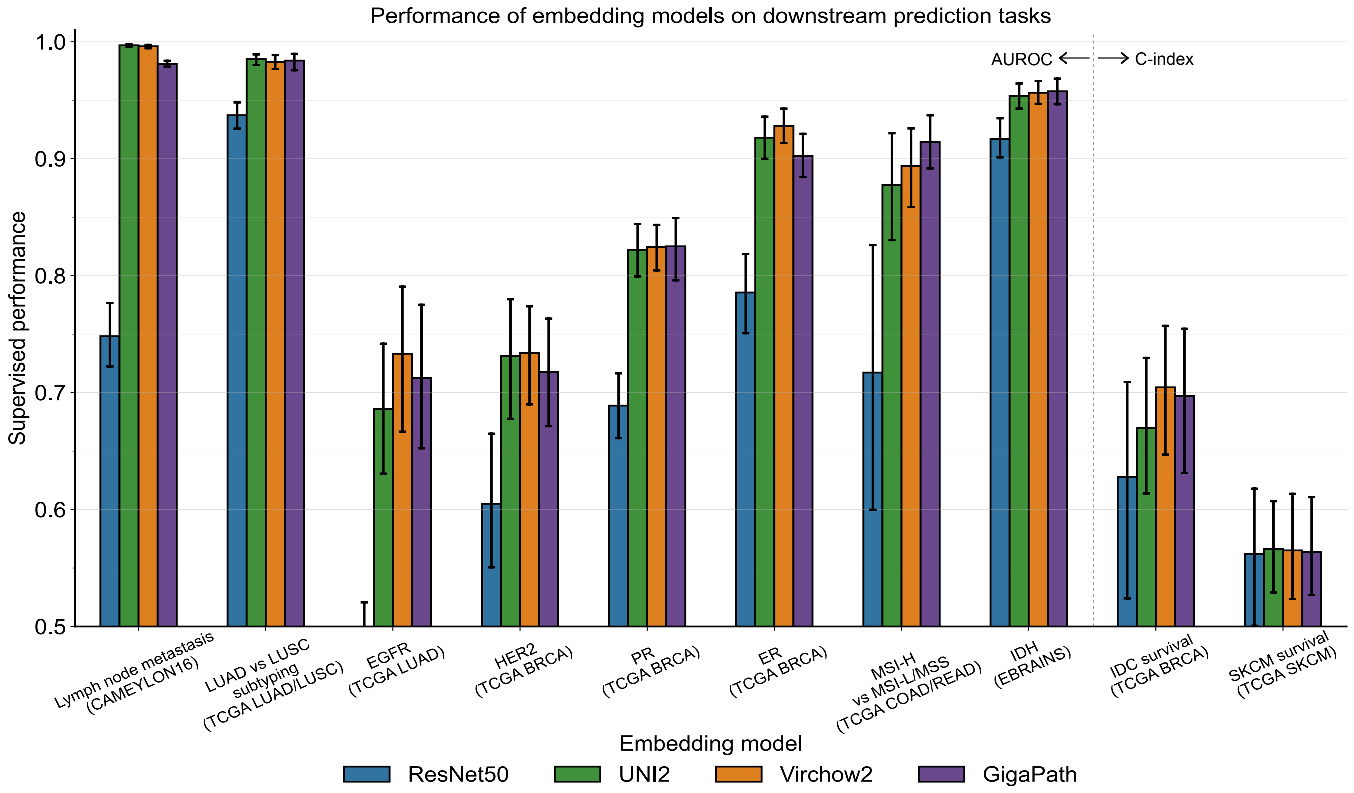


**Supplementary Fig. 1** | Performance of embedding models on downstream clinical tasks. Supervised performance is shown across ten diverse pathology tasks. The performance metric is the area under the receiver operating characteristic curve (AUROC) for classification tasks (left of the dotted line) and the concordance index (C-index) for survival tasks (right of the dotted line). Error bars represent 95% confidence intervals across five folds. The details of the tasks and data sources are described in Methods.

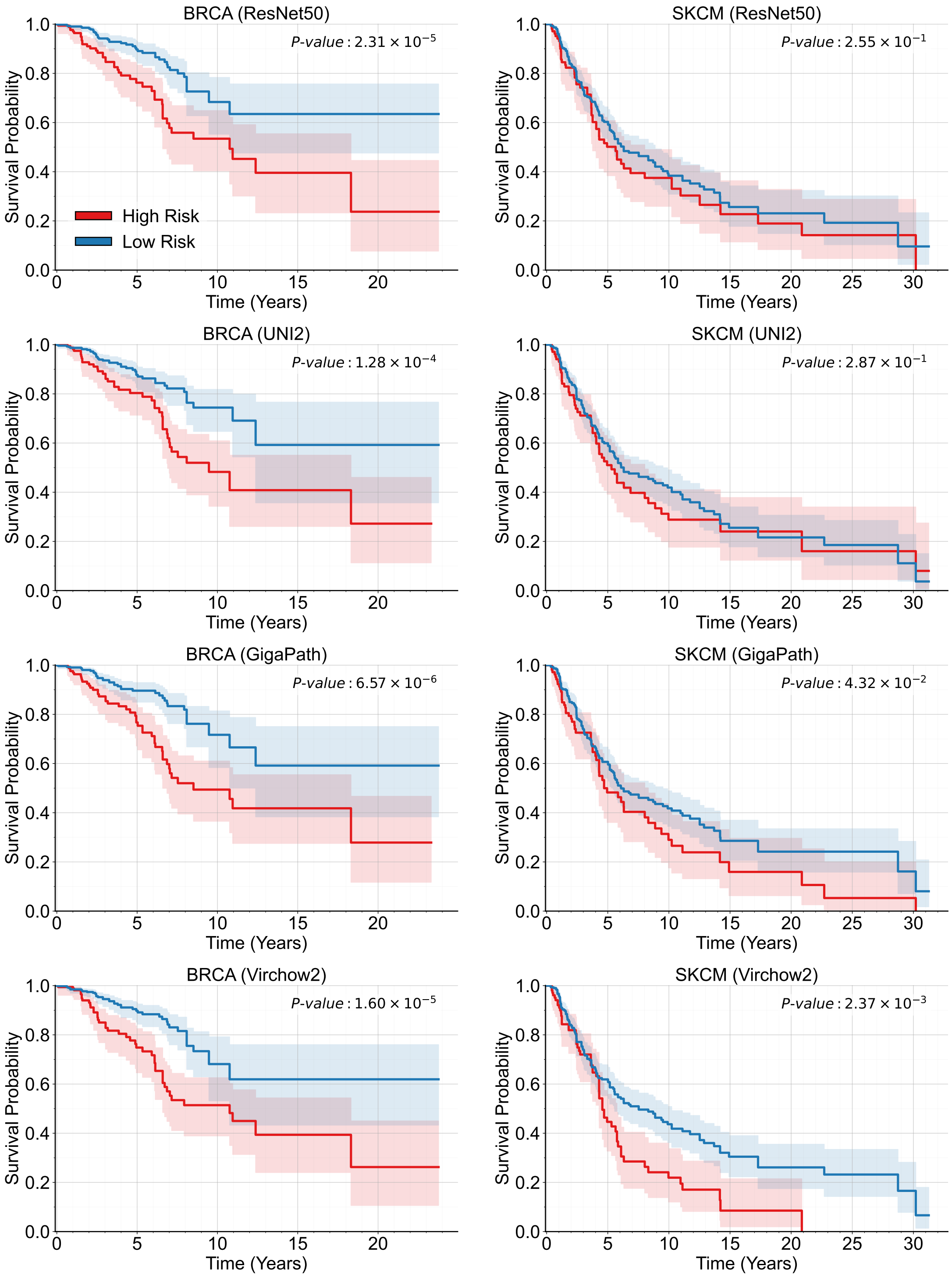


**Supplementary Fig. 2** | Performance of embedding models on survival prediction tasks.

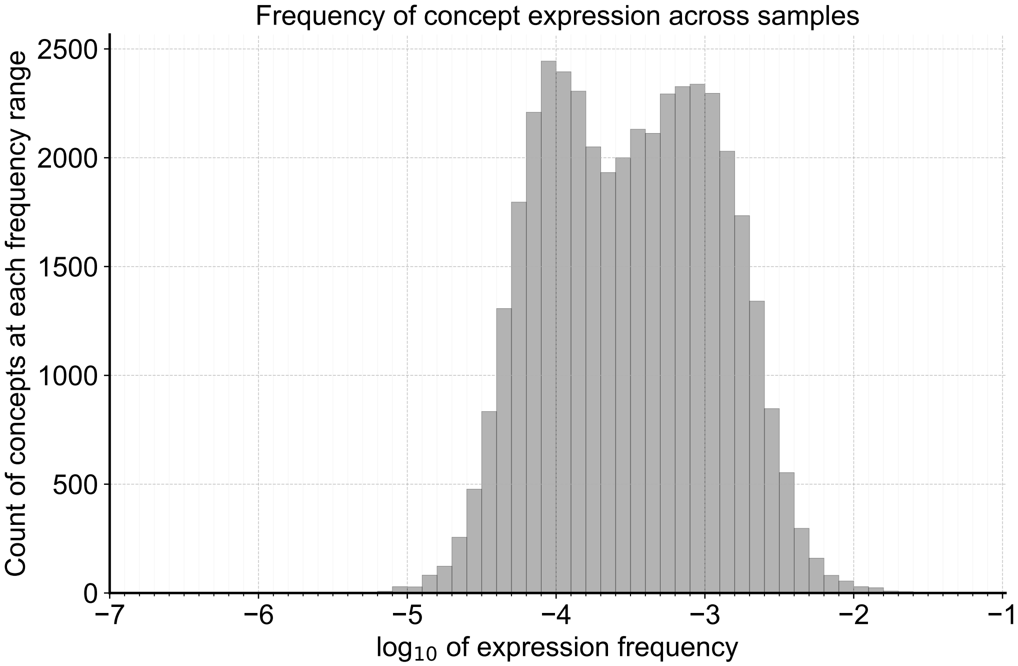


**Supplementary Fig. 3** | Frequency of concept nodes being activated. For each concept node in PICASSO, we computed how often it was activated across all images in the TCGA dataset. Among the 40,960 concept nodes, 79, 377, 3,400, 9,473 nodes showed activation frequencies higher than 1%, 0.5%, 0.2%, 0.1%, respectively.

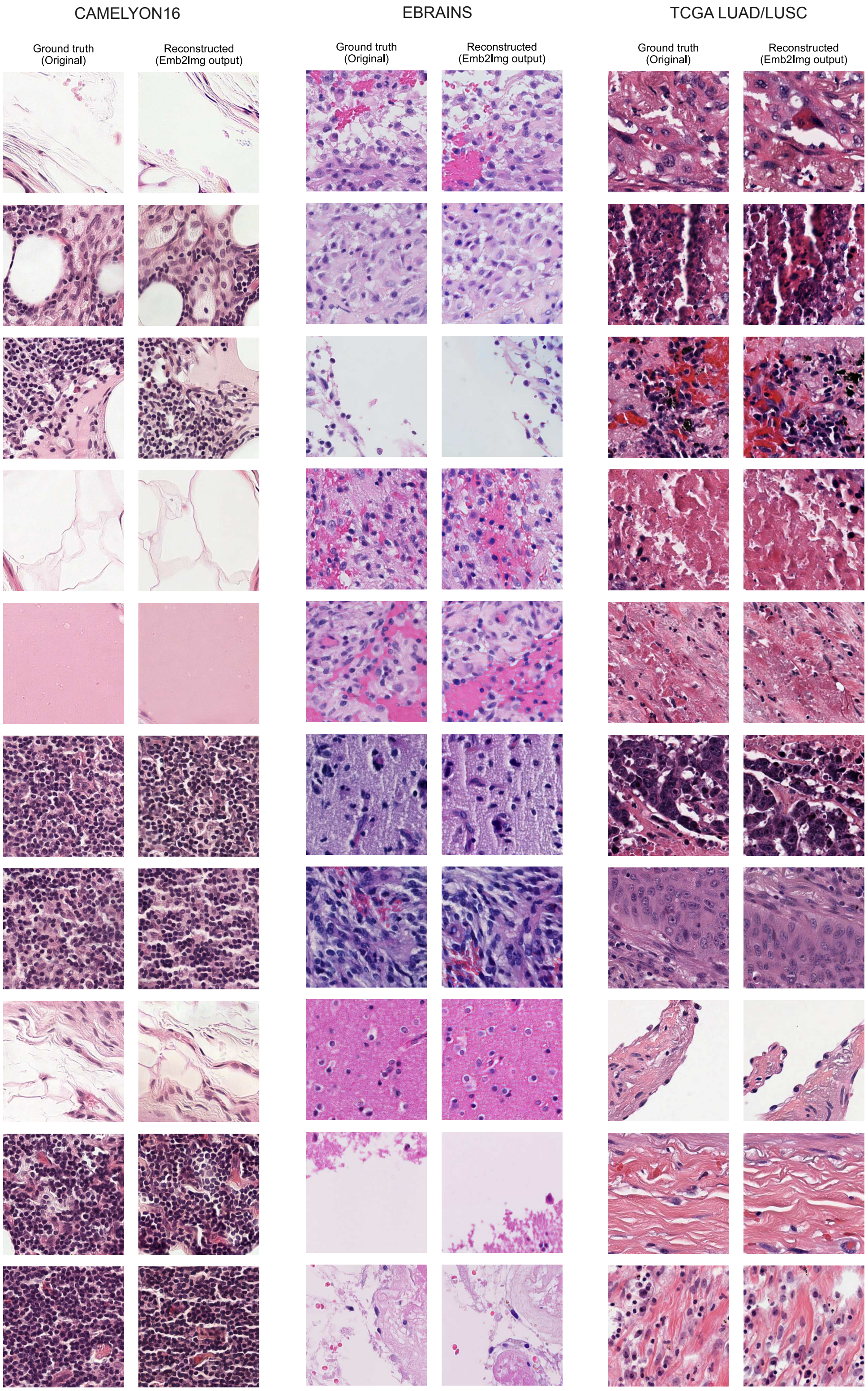


**Supplementary Fig. 4** | Performance of embedding models on downstream tasks.

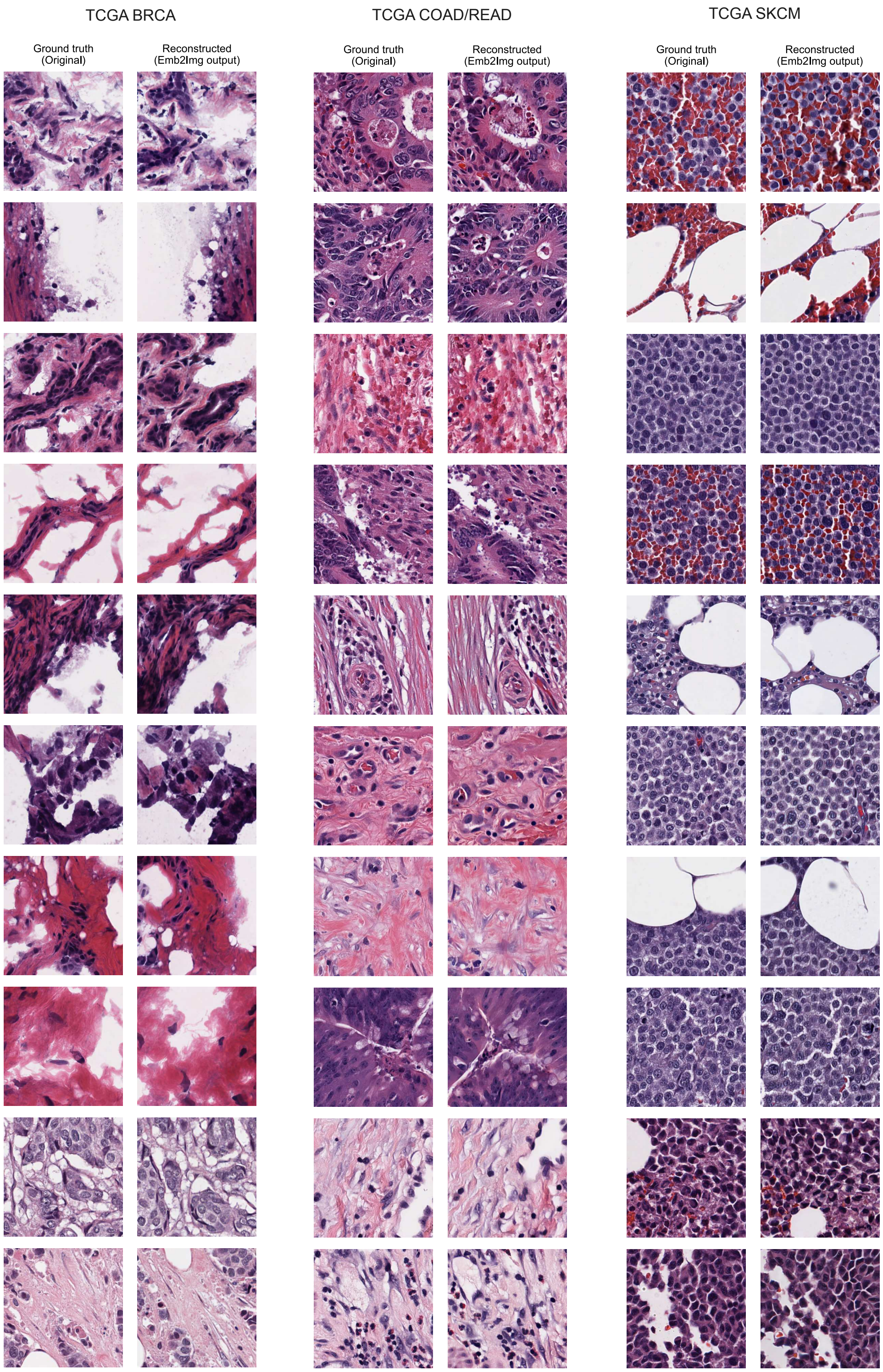


**Supplementary Fig. 5** | Performance of embedding models on downstream tasks.

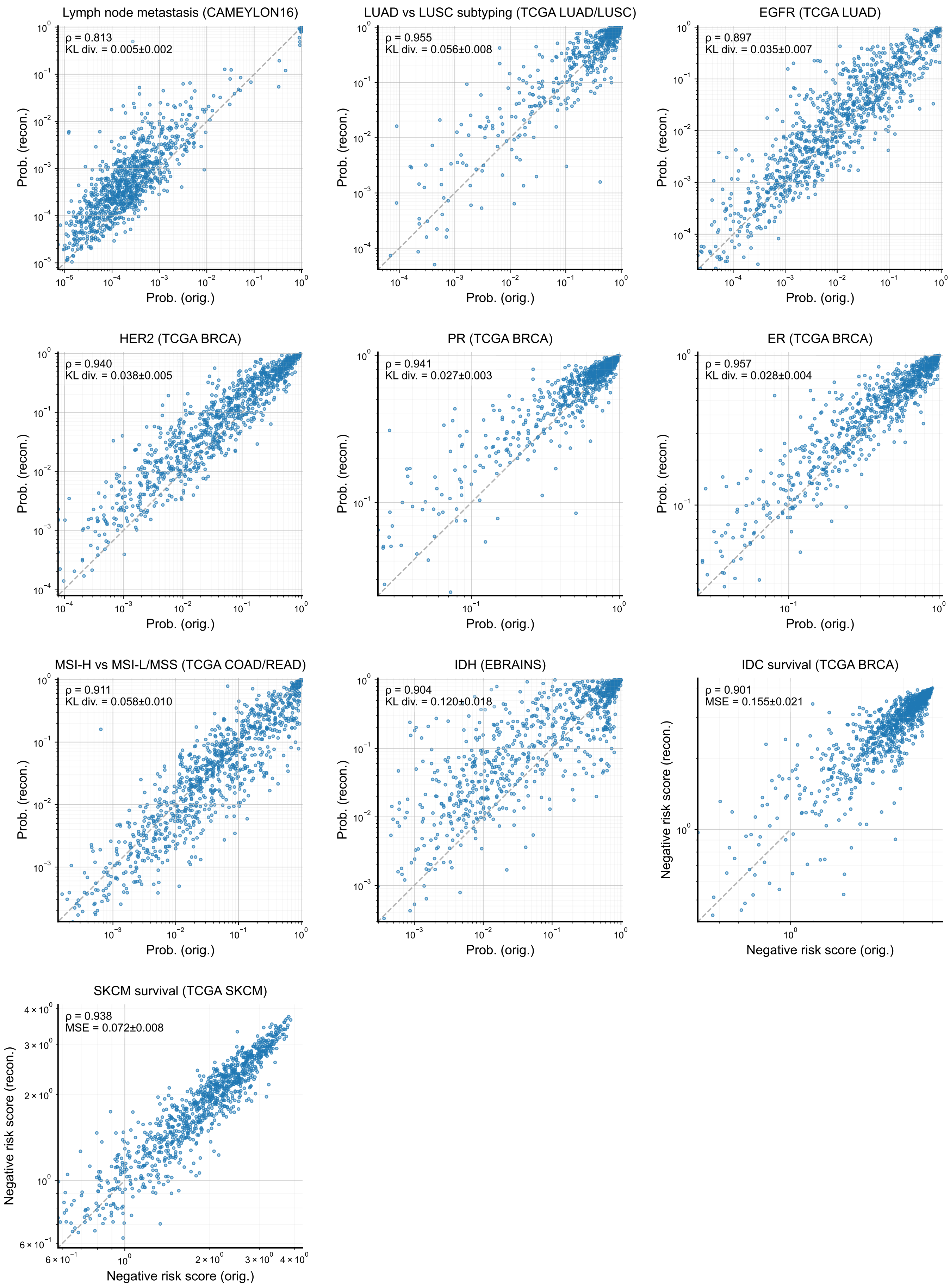


**Supplementary Fig. 6** | Performance of embedding models on downstream tasks.

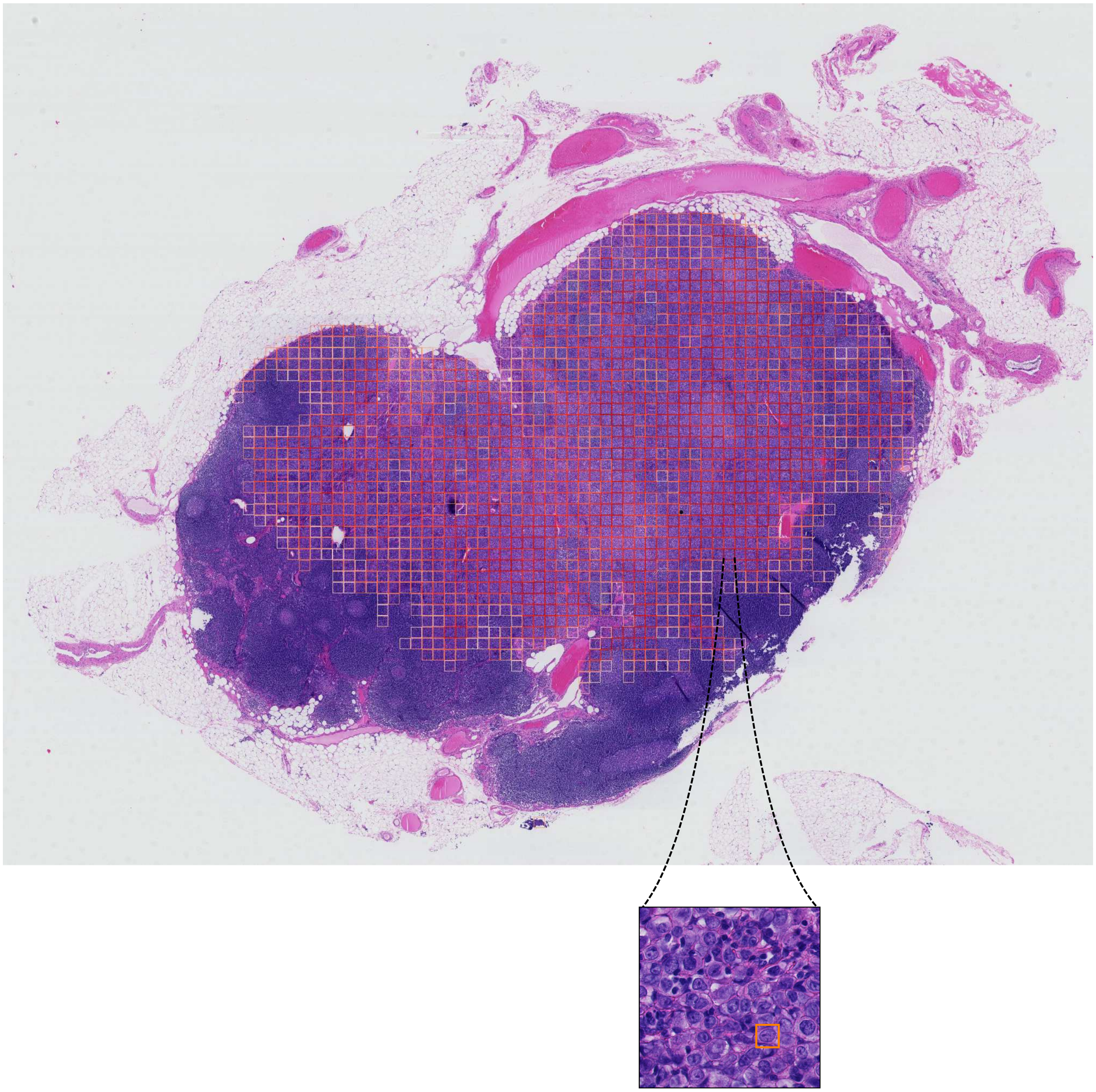

**Supplementary Fig. 7** | We highlight the regions that strongly express the first-ranked concept in the "test_114" slide from CAMELYON16, with color intensity reflecting expression strength.

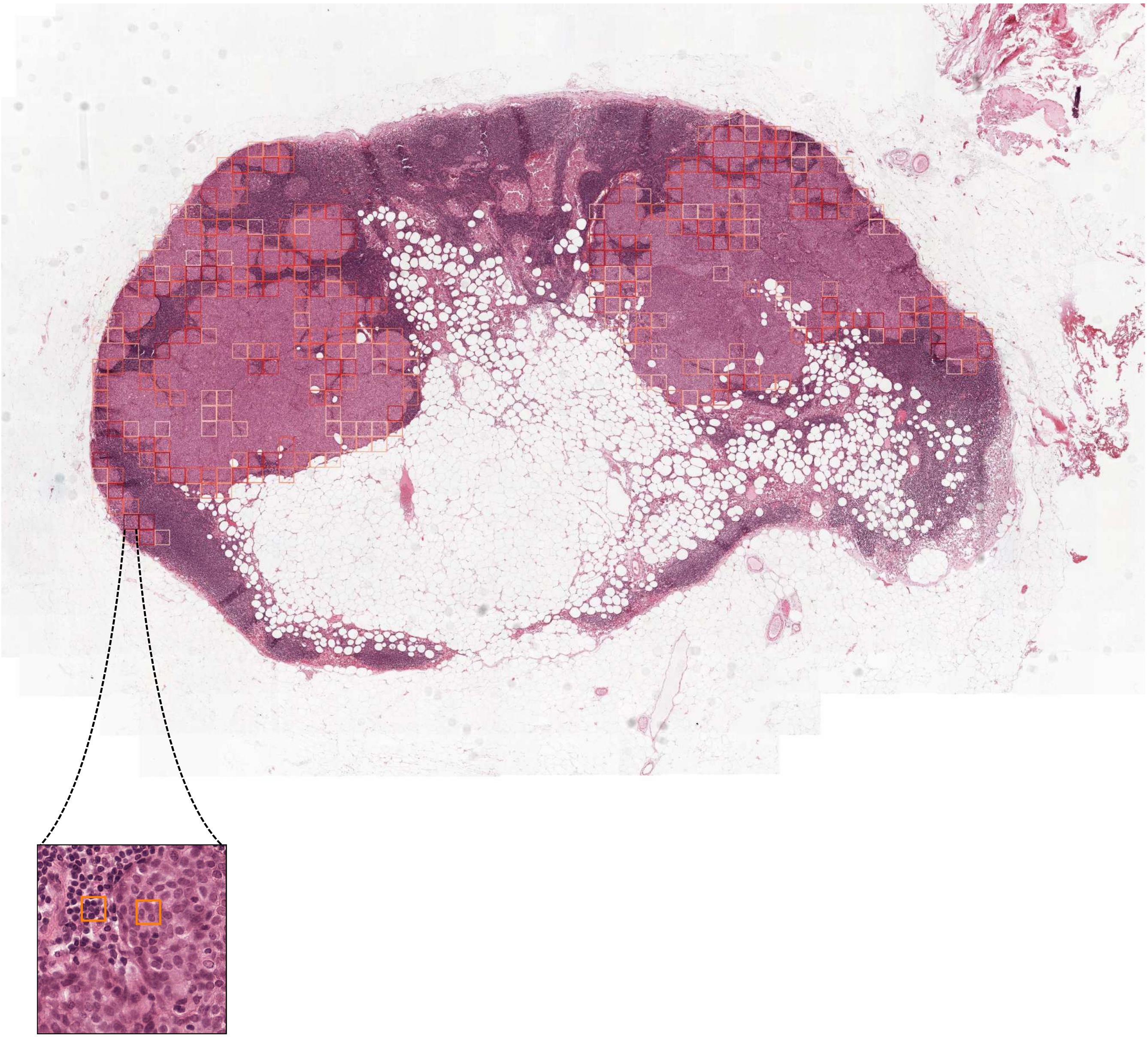

**Supplementary Fig. 8** | We highlight the regions that strongly express the second-ranked concept in the "test_040" slide from CAMELYON16, with color intensity reflecting expression strength.

Interpret the gene set:

PIGR, SNPH, C2orf72, S100A7, CES3, PROM1, ANKS1B, TMEM64, CCL28, GJC3, PIK3R1, ENPP3, DNAJC22, CYP4Z1, GPT, APOL4, IRAK3, CLGN, PSMA6, PKIA, EEF1A2, PDE10A, ZKSCAN1, SRP54, KMO, FAM177A1, ITGB6, AK4, JAKMIP1, USP1, PGM1, TACR1, TRIM4, KIAA0391, GABRE, DOCK7, GNA14, RBFOX2, C1orf198, EFHD1, PCDHB2, TXN2, NFIA, NPAS3, NREP, EIF3D

Provide 1. General overview, 2. Functional clusters, 3. Pathway-level insights, 4. Summary interpretation

**User**

**ChatGPT**

**1. General Overview**
The gene set looks heterogeneous, but there are recurring themes associated with:
• **Epithelial and secretory function**
• **Neural and synaptic signaling**
• **Metabolic and immune regulation**
These point to a mixture typical of tissue transcriptomes that include both epithelial and neuroendocrine or neuronal components (for example, airway epithelium, intestinal mucosa, or certain tumor types).

**2. Functional Clusters**
**A. Epithelial / Secretory surface markers**
• PIGR – Polymeric immunoglobulin receptor, mediates transcytosis of IgA/IgM in epithelial cells.
• S100A7 – Psoriasin, abundant in keratinocytes; involved in antimicrobial defense and skin inflammation.
• CCL28 – Chemokine expressed in mucosal epithelia; recruits IgA-secreting plasma cells.
• ITGB6 – Integrin subunit beta 6, often epithelial-specific, linked to TGF-β activation and wound repair.
• ENPP3, CES3 – Enzymes often expressed in glandular or secretory epithelia.
→ Suggests mucosal or epithelial barrier specialization.
**B. Immune / Inflammatory regulation**
• IRAK3 – Negative regulator of TLR signaling, dampens innate immune responses.
• APOL4 – Apolipoprotein, potential immune function.
• PSMA6, USP1 – Involved in proteasomal degradation and DNA damage/immune regulation.
→ Implies immune–epithelial signaling and stress response.
**C. Neural / Synaptic & signaling genes**
• SNPH (Syntaphilin) – Regulates synaptic vesicle transport in neurons.
• ANKS1B, GABRE, DOCK7, RBFOX2, NPAS3, NREP, KMO, PDE10A, JAKMIP1 – Highly enriched in neuronal tissue; roles in synapse formation, neurotransmitter signaling, and neurodevelopment.
→ Suggests neuronal or neuroendocrine identity, or cross-talk with neural pathways.
**D. Metabolism / Enzyme Activity**
• CYP4Z1, GPT, PGM1, AK4, PDE10A – Metabolic and signaling enzymes found in liver, brain, or certain carcinomas.
→ Indicates metabolic specialization, perhaps in energy metabolism or xenobiotic processing.
**E. Translational / protein synthesis and folding**
• EEF1A2, EIF3D, SRP54, CLGN – Components of translation machinery and protein folding.
→ Suggest high secretory or biosynthetic activity.

**3. Pathway-Level Insights**
If you were to run GO enrichment or Reactome analysis, you'd likely find:
• **Immune and epithelial defense processes**: mucosal immune response, cytokine signaling, epithelial barrier.
• **Neuron projection and synaptic processes**: regulation of neurotransmission, axon guidance.
• **Metabolism and oxidative stress**: oxidoreductase, energy metabolism.

**4. Summary Interpretation**
Overall, this gene set suggests a tissue or cell type characterized by:
• **Barrier/secretory epithelial features** (PIGR, S100A7, CCL28, ITGB6)
• **Strong neural or neuroendocrine signaling components** (SNPH, GABRE, NPAS3, RBFOX2, DOCK7)
• **Immune and metabolic modulation** (IRAK3, APOL4, CYP4Z1, GPT, AK4)

This could represent:
• A mucosal epithelial tissue interacting with immune and neural elements, or
• A neuroendocrine-like cell population with epithelial signatures, possibly in a developmental, inflamed, or tumor context.

**Supplementary Fig. 9** | Using a large language model to interpret the gene set associated with a visual concept. We prompted ChatGPT to interpret genes associated with the first-ranked concept in tumor detection (CAMELYON16). Only genes with Pearson correlation $> 0.2$ were included.

Interpret the gene set:

C2orf72, SNPH, ANKS1B, S100A7, PDE10A, JAKMIP1, MS4A1, EIF3L, DNAJC22, PGM1, PIK3R1, EEF1A2, CES3, CD37, ENPP3, PKIA, CYP4Z1, TMEM64, C1orf198, ABRACL, GJC3, RBFOX2, KIAA0391, TXN2, NFKBIA, IGHD, PPP1R1A, PSMA6, PPP2R3C, PIGR, FCRL1, USP1, EIF3D, ITGB6, DOCK7, SRP54, GNA14, EPHB4, UBE2E3

Provide 1. General overview, 2. Functional clusters, 3. Pathway-level insights, 4. Summary interpretation

**User**

**ChatGPT**

Let's break down this gene set step by step. I'll interpret this as if you provided a custom list from an enrichment or signature analysis, synthesizing biological themes and

**1. General Overview**
This gene set includes roughly 40 human protein-coding genes spanning:
• **Signal transduction and kinase activity** (e.g., PIK3R1, PDE10A, EPHB4, GNA14, PKIA, JAKMIP1)
• **Immune system and lymphocyte differentiation** (e.g., MS4A1, CD37, FCRL1, IGHD, PIGR, NFKBIA)
• **Protein synthesis and folding** (e.g., EIF3L, EIF3D, EEF1A2, DNAJC22, SRP54)
• **Metabolic enzymes** (e.g., PGM1, CES3, CYP4Z1)
• **Cell–cell communication and adhesion** (e.g., ITGB6, EPHB4, GJC3)
• **Regulatory/ubiquitin and phosphatase systems** (e.g., PPP1R1A, PPP2R3C, USP1, UBE2E3)
Overall, this looks like a mixed signature that combines immune regulatory, neuronal signaling, and metabolic/protein synthesis processes.

**2. Functional Clusters**
**A. Immune / B-cell–Associated Cluster**
• MS4A1 (CD20), CD37, FCRL1, IGHD, NFKBIA, PIGR → Characteristic of mature B lymphocytes and signaling molecules involved in B-cell receptor activation, antibody secretion, and NF-κB regulation.
**B. Signal Transduction / Kinase–Phosphatase Regulation**
• PIK3R1, PDE10A, PKIA, JAKMIP1, GNA14, EPHB4, TXN2, PPP1R1A, PPP2R3C → Reflects protein phosphorylation/dephosphorylation control, cAMP/cGMP balance, and growth/trophic signaling (PI3K, EPHB4, G protein pathways).
**C. Protein Synthesis, Folding, and Ubiquitin System**
• EIF3L, EIF3D, EEF1A2, SRP54, DNAJC22, USP1, UBE2E3 → Part of the translation initiation and protein quality control network, potentially marking a high biosynthetic state or stress response.
**D. Metabolic and Detoxification Enzymes**
• PGM1, CES3, CYP4Z1, ENPP3 → Enzymes involved in glucose metabolism, lipid/drug metabolism, and nucleotide metabolism, often upregulated in metabolically active or differentiating cells.
**E. Cell Adhesion / Developmental Signaling**
• ITGB6, EPHB4, GJC3, DOCK7, ANKS1B, SNPH → Links to cell migration, axonal transport, and tissue morphogenesis. Several (ANKS1B, SNPH) also show neural or synaptic associations.
**F. Other or Less Characterized Genes**
• C1orf198, C2orf72, KIAA0391, ABRACL, TMEM64 → Mostly uncharacterized or regulatory; may modulate mitochondrial function or cytoskeleton dynamics.

**3. Pathway-Level Insights**
Enrichment-based interpretation would likely highlight:
**A. PI3K–Akt signaling pathway**
• Core components: PIK3R1, PPP2R3C, PKIA, PPP1R1A, EPHB4, ITGB6
→ Suggesting cell survival, growth, and metabolic regulation themes.
**B. B-cell receptor signaling and NF-κB pathway**
• MS4A1, CD37, FCRL1, IGHD, NFKBIA, PIK3R1
→ Indicates activation, antigen presentation, or differentiation programs of B lymphocytes.
**C. Translation initiation / Ribosome pathway**
• EIF3D, EIF3L, EEF1A2, SRP54
→ Suggests translational upregulation or high protein output cells.
**D. Ubiquitin-mediated proteolysis & DNA damage control**
• USP1, UBE2E3, DNAJC22
→ Regulation of protein turnover and stress responses.
**E. Neuronal signaling / Cytoskeletal transport**
• SNPH, ANKS1B, DOCK7
→ If expression contexts are brain or neuron-specific datasets, these could reflect synaptic plasticity or axon maintenance.

**4. Summary Interpretation**
This gene set integrates multiple themes:
• **Immune lineage (especially B-cell) function**, reflected by MS4A1, CD37, FCRL1, and immune signaling scaffolds.
• **Signal transduction and kinase/phosphatase balance**, involving PIK3R1, PDE10A, EPHB4, linking immune and developmental pathways.
• **High biosynthetic activity**, with elevated translation and protein processing genes.
• **Metabolic reprogramming and protein turnover**, consistent with active differentiation or response to stimulation.
• **Potential neuronal or developmental links** (via ANKS1B, SNPH, DOCK7) suggesting cross-tissue expression or multifunctional pathways.

Overall, the gene list suggests a hybrid profile of activated B cells or secretory cells with intense signaling and biosynthetic activity, possibly indicating immune activation, cellular differentiation, or even cancerous transformation (e.g., B-cell lymphoma or highly active secretory tissue) depending on the biological context.

**Supplementary Fig. 10** | Using a large language model to interpret the gene set associated with a visual concept. We prompted ChatGPT to interpret genes associated with the second-ranked concept in tumor detection (CAMELYON16). Only genes with Pearson correlation $> 0.2$ were included.

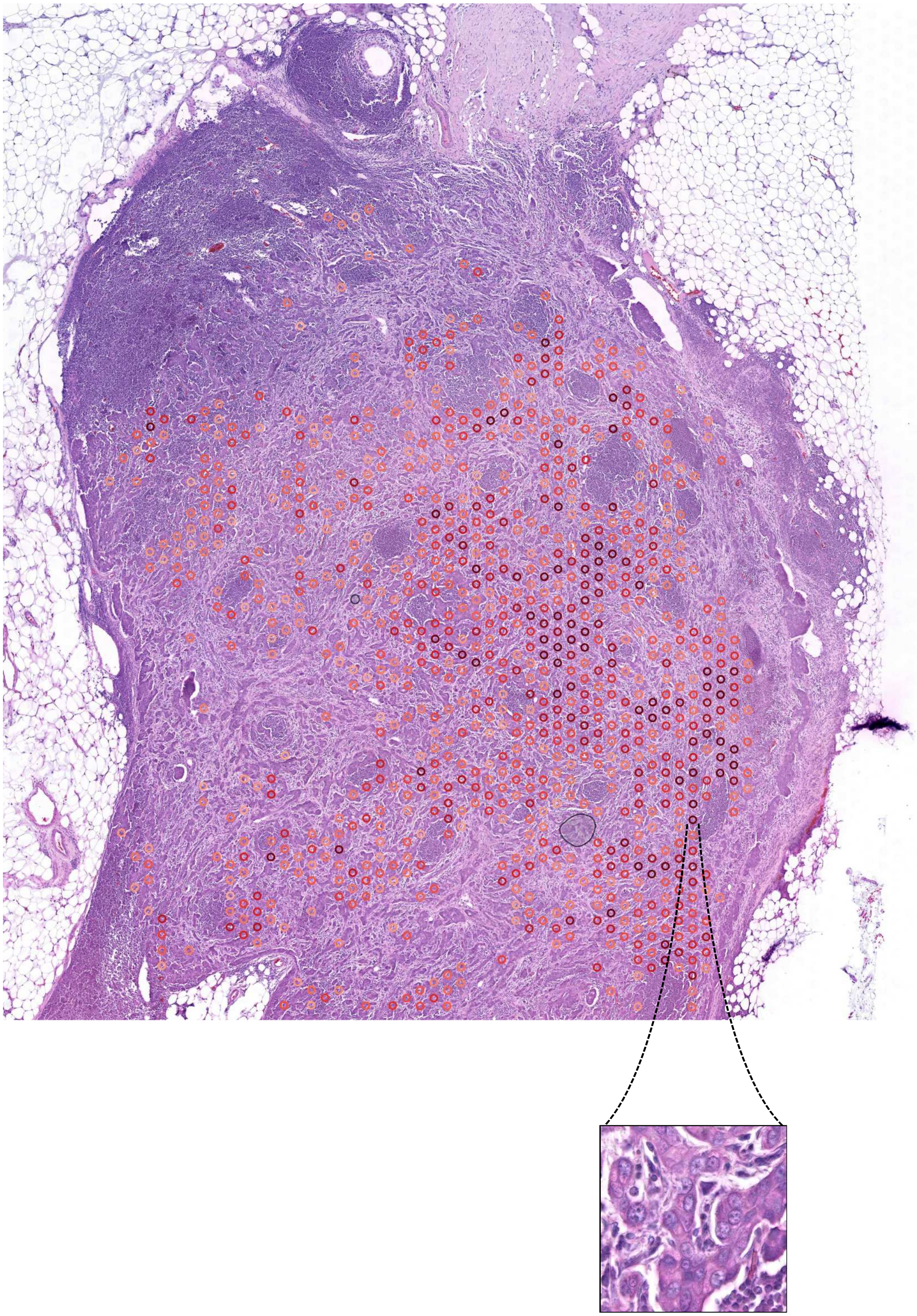

**Supplementary Fig. 11** | We highlight the regions that strongly express the first-ranked concept in the "NCBI684" spatial transcriptomics slide, with color intensity reflecting expression strength.

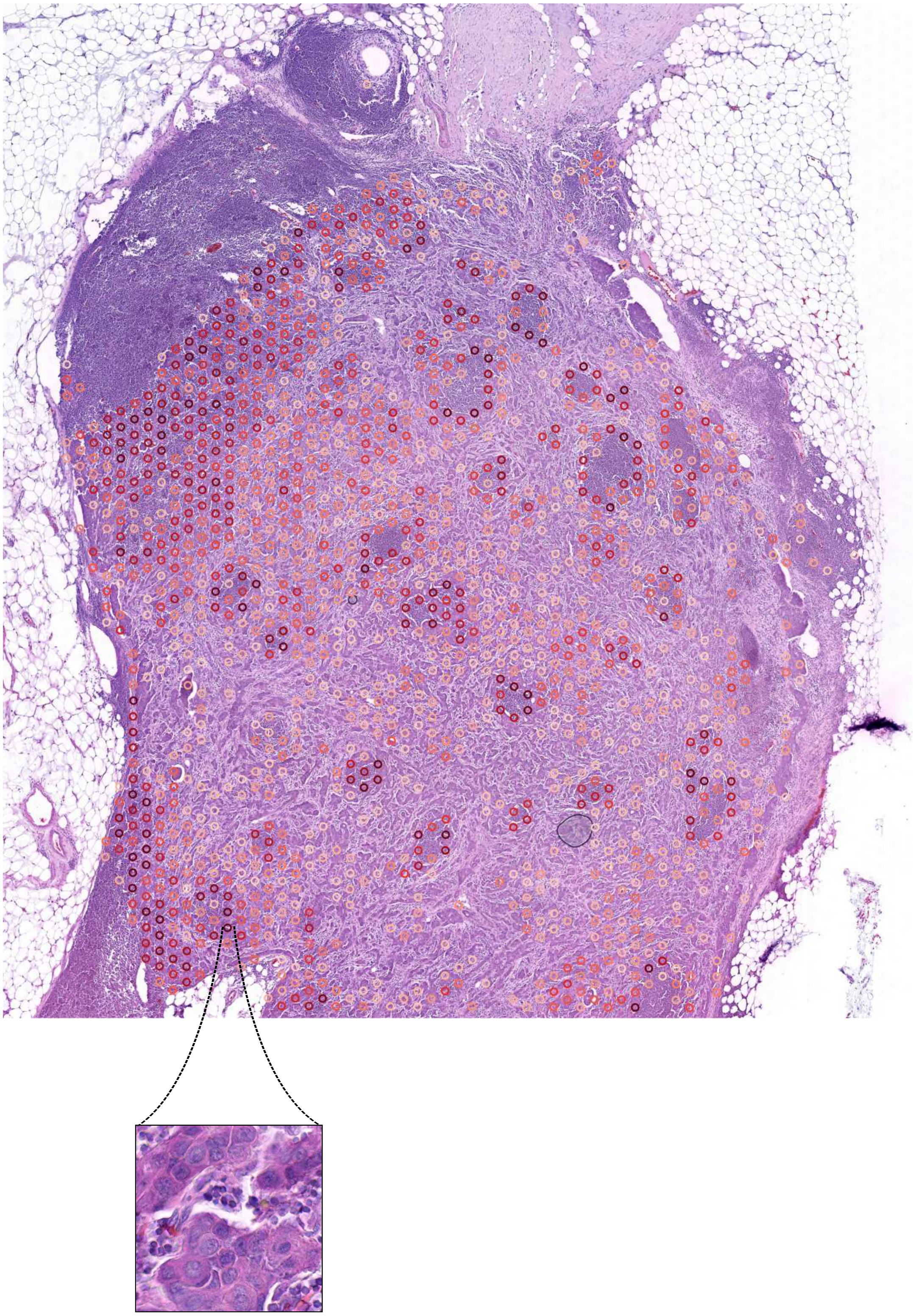

**Supplementary Fig. 12** | We highlight the regions that strongly express the second-ranked concept in the "NCBI684" spatial transcriptomics slide, with color intensity reflecting expression strength.

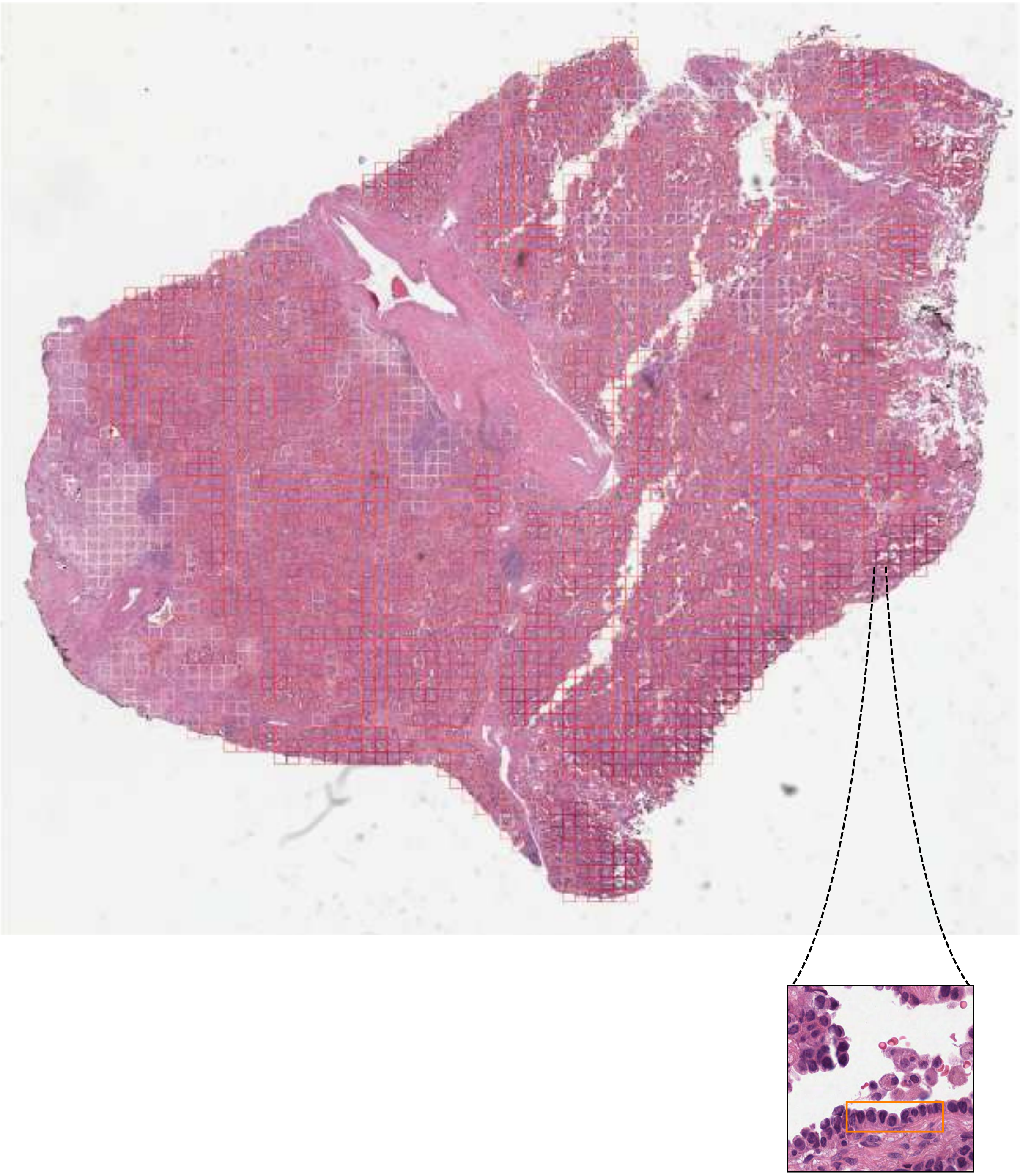

**Supplementary Fig. 13** | We highlight the regions that strongly express the first-ranked concept for the EGFR mutation model in the “TCGA-55-8087-01Z-00-DX1” slide from TCGA-LUAD, with color intensity reflecting expression strength.

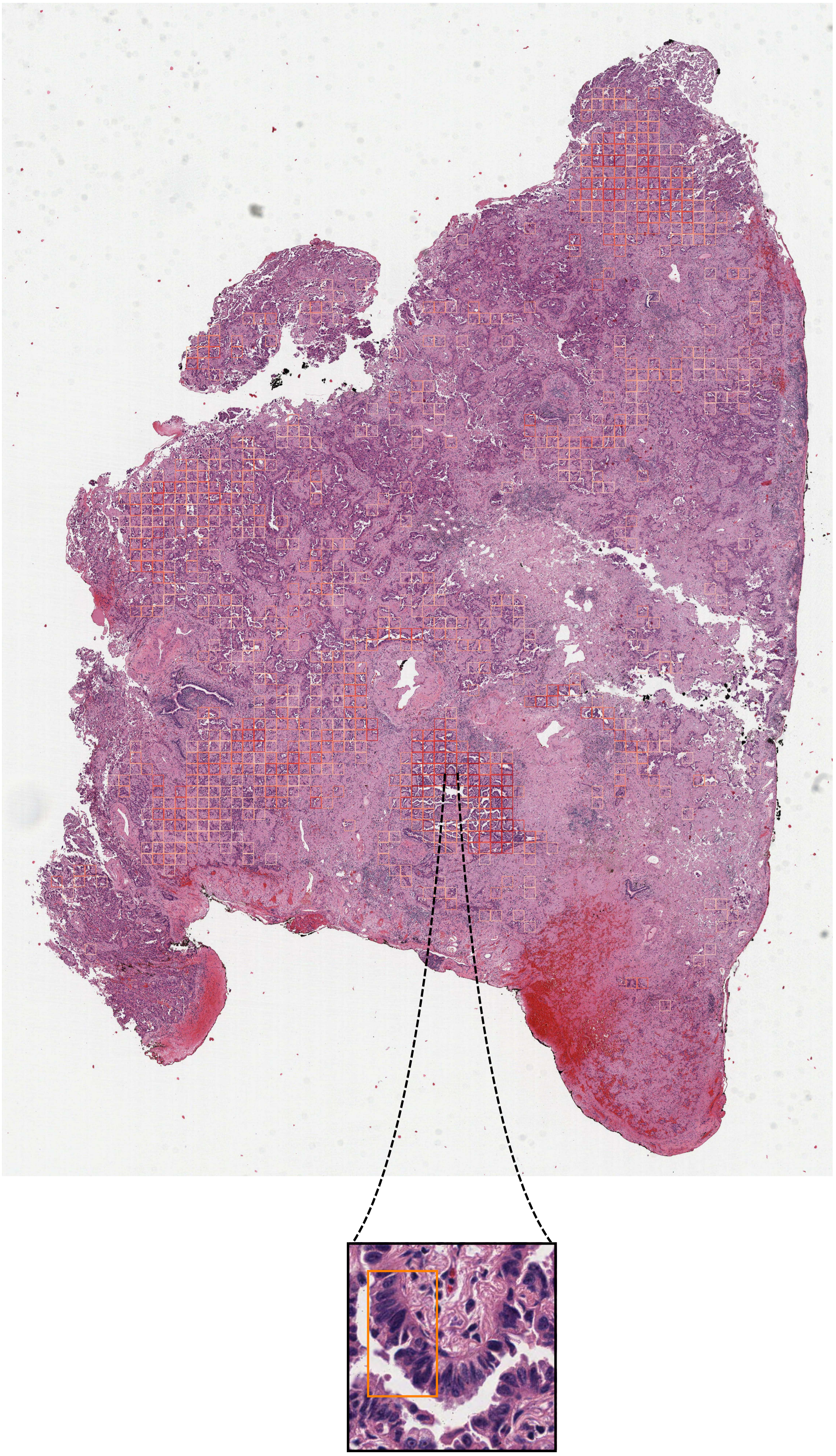

**Supplementary Fig. 14** | We highlight the regions that strongly express the second-ranked concept for the EGFR mutation model in the "TCGA-55-6980-01Z-00-DX1" slide from TCGA-LUAD, with color intensity reflecting expression strength.

Interpret the gene set:

CXCR4, MALL, CAPN8, PCSK2, PROX1, PTPRC, ACE2, GPX2, CCL5, CD14, GPR183, KLRB1, FCGR1A, VWA5A, TNFRSF17, AIF1, ESR1, FCER1A, STEAP4, TMC5, MZB1, LTBP2, SEMA3C, CFB, ANPEP, MS4A4A, SELL, PDGFRA, EGFR, CD1E, ADAM28, INMT, EGFL7, CD83, PLD4, CD79A, CD3E, CD1C, CD4, BASP1, MYLK, MS4A6A, CD3D, CYP2A7, PRDM1, CYTIP, DNAAF1, GZMK, CD28, APCDD1, CD1A, CCR7, CCR2, FOXP3, CLEC10A, CD247, CD27, VSIG4, CXCL2, SRPX, THY1, PECAM1, CTLA4

Provide 1. General overview, 2. Functional clusters, 3. Pathway-level insights, 4. Summary interpretation

**User**

**ChatGPT**

**1. General Overview**
This is a heterogeneous immune–stromal gene set containing a mix of:
• **Immune cell markers** (particularly T cells, B cells, monocytes/macrophages, and dendritic cells)
• **Endothelial and stromal components**
• **A few epithelial and signaling molecules** (ACE2, EGFR, PDGFRA, etc.)
Overall, it seems to represent a complex tissue microenvironment — likely immune infiltration within a stromal or epithelial tissue context, such as inflamed mucosa, tumor tissue, or lymphoid-rich organ.

**2. Functional Clusters**
Let's group by biological function or cell-type association:
**A. T cell–related genes**
• Core T-cell markers: CD3D, CD3E, CD247, CD28, CD4, GZMK, CCR7, CTLA4, FOXP3, CD27, CD2 (if present, but here CD27 serves similar role)
• Interpretation: Mix of $CD4^+$ helper (CD4, CCR7) and cytotoxic (GZMK) subsets; FOXP3 and CTLA4 indicate regulatory T cells (Tregs); CD27/CD28 are costimulatory receptors.
**B. B cell / plasma cell–related genes**
• Markers: CD79A, MZB1, TNFRSF17 (BCMA), PRDM1 (Blimp-1)
• Interpretation: Strong signatures of mature B cells and plasmablast/plasma cell differentiation.
**C. Monocyte-macrophage and dendritic cell genes**
• Markers: CD14, FCGR1A (FcγR1, CD64), FCER1A, AIF1 (Iba1), PTPRC (CD45), VSIG4, MS4A4A, CYTIP, CD83, CD1A, CD1C, CD1E, CLEC10A, CCR2
• Interpretation: Classical and non-classical monocyte/macrophage markers; antigen-presenting dendritic cells (particularly $CD1^+$ subsets); immunomodulatory macrophages (VSIG4).
**D. Chemokines and chemokine receptors**
• CXCR4, CCR7, CCR2, CCL5 (RANTES), CXCL2, GPR183 (EBI2)
• These specify immune cell trafficking, positioning, and inflammation.
  • CCR7 → lymphoid homing
  • CCL5/CXCL2 → recruitment of effector T cells & neutrophils
  • CXCR4 → retention in niches (e.g., bone marrow, Tfh zones)
**E. Endothelial, stromal, and structural genes**
• PECAM1 (CD31), PDGFRA, EGFR, THY1, LTBP2, SEMA3C, EGFL7, VWA5A, MALL
• Indicate endothelial, fibroblast, and tissue structural components.
These are likely vascular and matrix-associated signals — suggesting immune infiltration within an endothelial/stromal compartment.
**F. Enzymes and metabolic/stress response**
• ACE2 (angiotensin-converting enzyme 2), GPX2 (glutathione peroxidase 2), CAPN8, PCSK2, ANPEP (CD13), STEAP4, INMT, MYLK, CYP2A7, PLD4
• Implicate metabolic regulation, peptide processing, redox balance, or proteolysis within tissue or immune cells.
**G. Signaling / transcriptional regulators**
• PROX1 – developmental transcription factor (lymphangiogenesis, vascular identity)
• ESR1 – nuclear hormone receptor (estrogen receptor alpha)
• APCDD1 – Wnt pathway modulator
• FOXP3, PRDM1 – transcription factors for Treg and plasma cell differentiation.
• These suggest the tissue context could be influenced by hormonal, developmental, or immune differentiation signals.

**3. Pathway-Level Insights**
**A. Immune System Pathways**
Adaptive immune activation:
• Antigen presentation (CD1A/C/D/E, HLA not listed but context implied)
• TCR signaling (CD3 complex, CD28/CD27 costimulation)
• BCR/plasma cell differentiation (CD79A, PRDM1, TNFRSF17)
Immune regulation:
FOXP3, CTLA4, VSIG4 → immunosuppressive networks and checkpoint control
Chemotaxis and cytokine signaling:
CCL5, CXCL2, CCR2, CXCR4 → recruitment and positioning
Myeloid activation & interferon response:
FCGR1A, AIF1, FCER1A, CFB → innate inflammatory and complement pathways
**B. Stromal/Vascular Pathways**
Angiogenesis & endothelial regulation:
PECAM1, EGFL7, PROX1, PDGFRA, THY1
Extracellular matrix remodeling:
LTBP2, SEMA3C, ADAM28
Barrier & epithelial signaling:
ACE2, EGFR, ANPEP, GPX2

**4. Summary Interpretation**
The gene set indicates a coordinated immune-stromal activation state, integrating adaptive immune cell activity (T, B, and antigen-presenting cells) with vascular and matrix remodeling responses — characteristic of a tissue undergoing immune infiltration, inflammation, or immune-driven transformation.

**Supplementary Fig. 15** | Using a large language model to interpret the gene set associated with a visual concept. We prompted ChatGPT to interpret genes associated with the first-ranked concept in the EGFR mutation status prediction model. Only genes with Pearson correlation > 0.3 were included.

**User**

Interpret the gene set:

ACE2, PROX1, PCSK2, GPX2, MALL, CAPN8, CXCR4, STEAP4, TNFRSF17, FCGR1A, ANPEP, ESR1, VWA5A, TMC5, EGFL7, INMT, EGFR, PTPRC, CYP2A7, MZB1, CD83, CXCL2, FCER1A, DNAAF1, CD1E, GPR183, SEMA3C, SELL, LTBP2, CCL5, CD1C, CD14, PLD4, PDGFRA, KLRB1, ADAM28, CD79A, S100A1, SOX2, MYLK, AIF1, SNTN, APCDD1, CD1A

Provide 1. General overview, 2. Functional clusters, 3. Pathway-level insights, 4. Summary interpretation

**ChatGPT**

**1. General Overview**
This gene set combines epithelial, immune, and stromal/vascular markers, indicating a complex tissue environment where epithelial cells interact with immune and supporting stromal elements.

**2. Functional Clusters**
**A. Immune-related genes:** CD1A/C/E, CD14, CD79A, CD83, FCGR1A, TNFRSF17, CXCR4, CCL5, CXCL2, KLRB1, MZB1, GPR183, PLD4, PTPRC.
→ Indicate B cells, monocytes/macrophages, dendritic cells, and active cytokine/chemokine signaling.
**B. Epithelial/progenitor genes:** ACE2, ANPEP, GPX2, EGFR, SOX2, PROX1, PCSK2, CAPN8, MALL.
→ Reflect epithelial barrier function, progenitor activity, and surface enzyme expression.
**C. Stromal/vascular genes:** EGFL7, PDGFRA, LTBP2, STEAP4, VWA5A, APCDD1.
→ Suggest endothelial remodeling, fibroblast signaling, and ECM organization.
**D. Regulatory/metabolic genes:** ESR1, MYLK, ADAM28, S100A1, INMT.
→ Link hormonal, mechanical, and metabolic modulation.

**3. Pathway-Level Insights**
Based on known associations:
**Immune activation pathways:**
• Antigen processing & presentation (CD1 family, FC receptors)
• B-cell receptor signaling (CD79A, TNFRSF17, MZB1)
• Chemokine-mediated leukocyte migration (CXCR4, CCL5, CXCL2)
**Barrier and epithelial signaling:**
• EGF/EGFR axis and epithelial differentiation (EGFR, PROX1, SOX2)
• Protease activity and peptide processing (ANPEP, ACE2, PCSK2, CAPN8)
• Oxidative stress defenses (GPX2)
**Vascular and extracellular environment:**
• ECM organization, angiogenesis, PDGF and TGF-β signaling (PDGFRA, LTBP2, EGFL7)
**Crosstalk:**
The gene combination suggests crosstalk between epithelial barrier cells and immune cells through chemokine signaling (CXCL2, CCL5, CXCR4) and possibly tissue remodeling/repair pathways mediated by fibroblasts and endothelial cells.

**4. Summary Interpretation**
This gene set represents a multicellular tissue environment characterized by epithelial–immune–stromal interactions, consistent with:
• **An inflamed or regenerating epithelial tissue** (with epithelial markers ACE2, EGFR, SOX2, ANPEP, and immune infiltration signatures).
• **Active immune compartment** including B cells, dendritic cells, and monocytes (CD1, CD14, CD83, PTPRC, CD79A).
• **Stromal and vascular remodeling** indicating microenvironmental adaptation (EGFL7, PDGFRA, LTBP2).
• **Possible hormonal and metabolic modulatory influences** (ESR1, STEAP4, MYLK, ADAM28).
Overall, these patterns are consistent with a differentiated epithelial tissue undergoing repair, inflammatory response, or immune surveillance, such as might be observed in mucosal surfaces (e.g., respiratory or intestinal epithelium) or tumor-adjacent tissue with immune cell infiltration.

**Optional summary phrase:**
"Integrated interpretation: The gene set points to epithelial barrier cells interacting with infiltrating B cells, antigen-presenting cells, and stromal elements—driven by cytokine/chemokine signaling, growth factor pathways (EGFR/PDGF), and oxidative/metabolic regulation (GPX2/STEAP4)—indicative of a tissue remodeling or immune-activated state."

**Supplementary Fig. 16** | Using a large language model to interpret the gene set associated with a visual concept. We prompted ChatGPT to interpret genes associated with the second-ranked concept in the EGFR mutation status prediction model. Only genes with Pearson correlation $> 0.3$ were included.

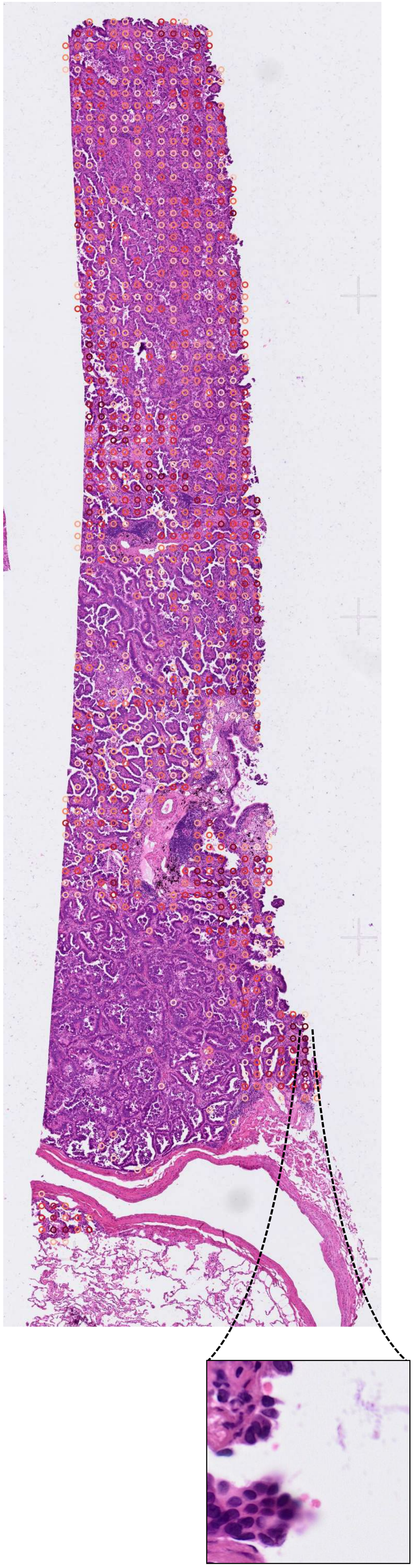

**Supplementary Fig. 17** | We highlight the regions that strongly express the first-ranked concept in the "TENX118" spatial transcriptomics slide, with color intensity reflecting expression strength.

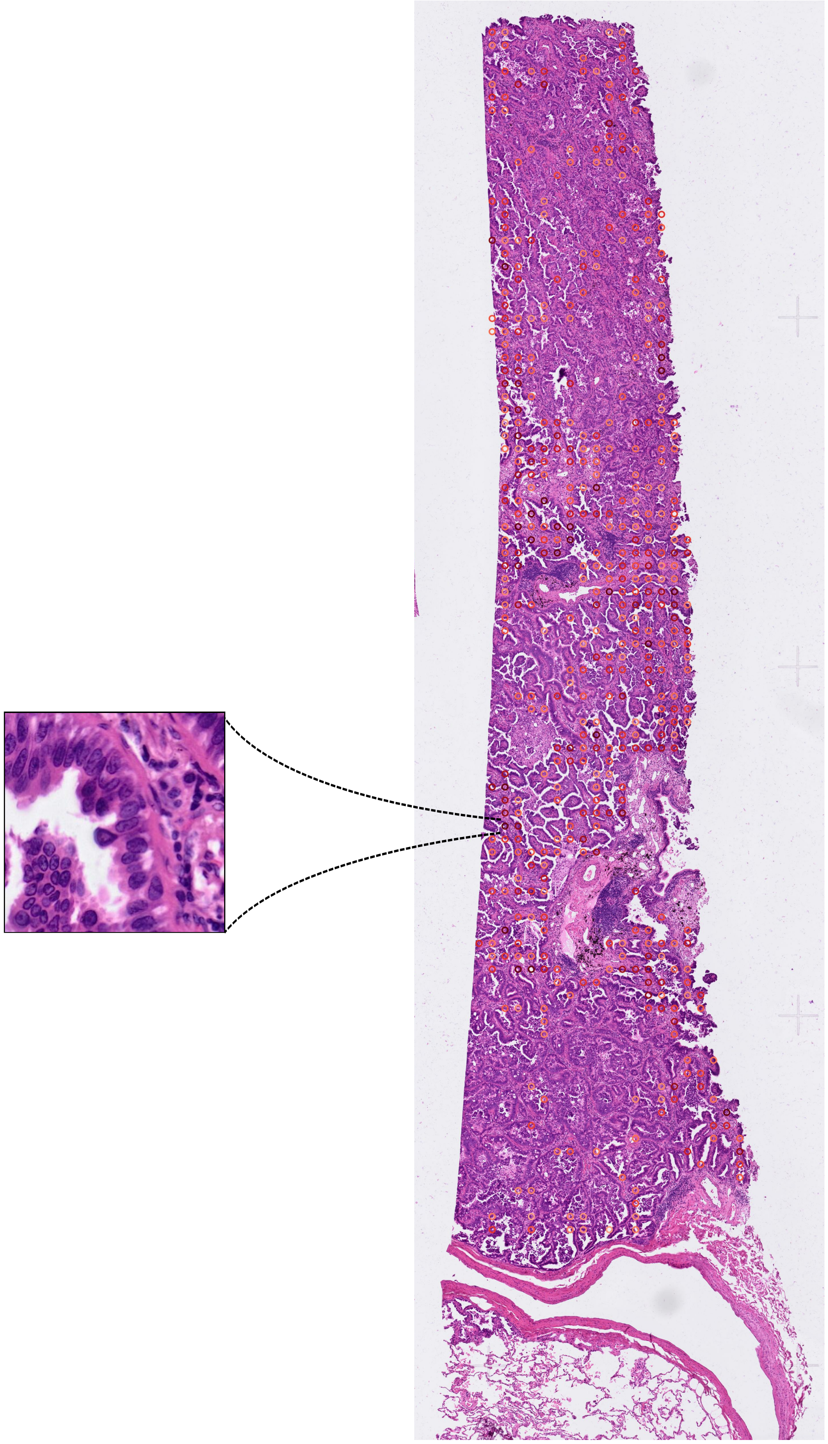

**Supplementary Fig. 18** | We highlight the regions that strongly express the second-ranked concept in the "TENX118" spatial transcriptomics slide, with color intensity reflecting expression strength.

**Supplementary Table 1** | Sample counts of TCGA cancer types

| **Cancer type** | **Abbreviation** | **Sample count** |
|---|---|---:|
| Breast invasive carcinoma | BRCA | 1,121 |
| Low grade glioma | LGG | 838 |
| Glioblastoma multiforme | GBM | 817 |
| Sarcoma | SARC | 600 |
| Uterine corpus endometrial carcinoma | UCEC | 566 |
| Lung adenocarcinoma | LUAD | 530 |
| Kidney renal clear cell carcinoma | KIRC | 519 |
| Thyroid carcinoma | THCA | 518 |
| Lung squamous cell carcinoma | LUSC | 512 |
| Skin cutaneous melanoma | SKCM | 473 |
| Head and Neck squamous cell carcinoma | HNSC | 472 |
| Bladder urothelial carcinoma | BLCA | 457 |
| Prostate adenocarcinoma | PRAD | 449 |
| Colon adenocarcinoma | COAD | 441 |
| Stomach adenocarcinoma | STAD | 399 |
| Liver hepatocellular carcinoma | LIHC | 371 |
| Kidney renal papillary cell carcinoma | KIRP | 297 |
| Cervical & endocervical cancer | CESC | 279 |
| Testicular germ cell tumors | TGCT | 254 |
| Adrenocortical carcinoma | ACC | 227 |
| Pancreatic adenocarcinoma | PAAD | 209 |
| Pheochromocytoma and Paraganglioma | PCPG | 195 |
| Thymoma | THYM | 180 |
| Rectum adenocarcinoma | READ | 158 |
| Esophageal carcinoma | ESCA | 158 |
| Kidney chromophobe | KICH | 121 |
| Ovarian serous cystadenocarcinoma | OV | 107 |
| Uterine carcinosarcoma | UCS | 87 |
| Mesothelioma | MESO | 86 |
| Uveal melanoma | UVM | 80 |
| Diffuse large B-cell lymphoma | DLBC | 43 |
| Cholangiocarcinoma | CHOL | 39 |
| **Total** | | **11,603** |

**Supplementary Table 2** | Gene association results corresponding to the first-ranked concept in tumor detection (CAMELYON16). Genes with Pearson's $r > 0.3$ are displayed.

| **Gene** | **Correlation** |
|---|---|
| PIGR | 0.491 |
| SNPH | 0.449 |
| C2orf72 | 0.422 |
| S100A7 | 0.407 |
| CES3 | 0.396 |
| PROM1 | 0.394 |
| ANKS1B | 0.390 |
| TMEM64 | 0.388 |
| CCL28 | 0.386 |
| GJC3 | 0.385 |
| PIK3R1 | 0.377 |
| ENPP3 | 0.372 |
| DNAJC22 | 0.368 |
| CYP4Z1 | 0.367 |
| GPT | 0.366 |
| APOL4 | 0.363 |
| IRAK3 | 0.360 |
| CLGN | 0.354 |
| PSMA6 | 0.352 |
| PKIA | 0.348 |
| EEF1A2 | 0.345 |
| PDE10A | 0.345 |
| ZKSCAN1 | 0.341 |
| SRP54 | 0.339 |
| KMO | 0.338 |
| FAM177A1 | 0.336 |
| ITGB6 | 0.335 |
| AK4 | 0.335 |
| JAKMIP1 | 0.329 |
| USP1 | 0.328 |
| PGM1 | 0.324 |
| TACR1 | 0.323 |
| TRIM4 | 0.322 |
| KIAA0391 | 0.322 |
| GABRE | 0.322 |
| DOCK7 | 0.322 |
| GNA14 | 0.316 |
| RBFOX2 | 0.315 |
| C1orf198 | 0.314 |
| EFHD1 | 0.310 |
| PCDHB2 | 0.310 |
| TXN2 | 0.309 |
| NFIA | 0.307 |
| NPAS3 | 0.306 |
| NREP | 0.305 |
| EIF3D | 0.304 |

**Supplementary Table 3** | Gene association results corresponding to the second-ranked concept in tumor detection (CAMELYON16). Genes with Pearson's $r > 0.3$ are displayed.

| **Gene** | **Correlation** |
|---|---:|
| C2orf72 | 0.368 |
| SNPH | 0.358 |
| ANKS1B | 0.356 |
| S100A7 | 0.355 |
| PDE10A | 0.353 |
| JAKMIP1 | 0.349 |
| MS4A1 | 0.344 |
| EIF3L | 0.344 |
| DNAJC22 | 0.343 |
| PGM1 | 0.341 |
| PIK3R1 | 0.340 |
| EEF1A2 | 0.339 |
| CES3 | 0.339 |
| CD37 | 0.336 |
| ENPP3 | 0.334 |
| PKIA | 0.331 |
| CYP4Z1 | 0.330 |
| TMEM64 | 0.329 |
| C1orf198 | 0.329 |
| ABRACL | 0.329 |
| GJC3 | 0.327 |
| RBFOX2 | 0.327 |
| KIAA0391 | 0.326 |
| TXN2 | 0.326 |
| NFKBIA | 0.324 |
| IGHD | 0.324 |
| PPP1R1A | 0.324 |
| PSMA6 | 0.322 |
| PPP2R3C | 0.320 |
| PIGR | 0.319 |
| FCRL1 | 0.318 |
| USP1 | 0.318 |
| EIF3D | 0.314 |
| ITGB6 | 0.313 |
| DOCK7 | 0.308 |
| SRP54 | 0.306 |
| GNA14 | 0.306 |
| EPHB4 | 0.305 |
| UBE2E3 | 0.304 |

**Supplementary Table 4** | Gene association results corresponding to the first-ranked concept in EGFR predictions (TCGA-LUAD). Genes with Pearson's $r > 0.2$ are displayed.

| Gene | Correlation | Gene | Correlation |
|---|---|---|---|
| CXCR4 | 0.453 | MYLK | 0.255 |
| MALL | 0.439 | MS4A6A | 0.250 |
| CAPN8 | 0.394 | CD3D | 0.248 |
| PCSK2 | 0.394 | CYP2A7 | 0.246 |
| PROX1 | 0.391 | PRDM1 | 0.245 |
| PTPRC | 0.387 | CYTIP | 0.242 |
| ACE2 | 0.384 | DNAAF1 | 0.231 |
| GPX2 | 0.377 | GZMK | 0.231 |
| CCL5 | 0.369 | CD28 | 0.227 |
| CD14 | 0.363 | APCDD1 | 0.225 |
| GPR183 | 0.351 | CD1A | 0.223 |
| KLRB1 | 0.350 | CCR7 | 0.222 |
| FCGR1A | 0.342 | CCR2 | 0.222 |
| VWA5A | 0.338 | FOXP3 | 0.220 |
| TNFRSF17 | 0.325 | CLEC10A | 0.217 |
| AIF1 | 0.324 | CD247 | 0.215 |
| ESR1 | 0.316 | CD27 | 0.213 |
| FCER1A | 0.307 | VSIG4 | 0.211 |
| STEAP4 | 0.302 | CXCL2 | 0.207 |
| TMC5 | 0.297 | SRPX | 0.207 |
| MZB1 | 0.294 | THY1 | 0.203 |
| LTBP2 | 0.294 | PECAM1 | 0.203 |
| SEMA3C | 0.293 | CTLA4 | 0.201 |
| CFB | 0.293 | | |
| ANPEP | 0.292 | | |
| MS4A4A | 0.291 | | |
| SELL | 0.291 | | |
| PDGFRA | 0.288 | | |
| EGFR | 0.288 | | |
| CD1E | 0.286 | | |
| ADAM28 | 0.285 | | |
| INMT | 0.285 | | |
| EGFL7 | 0.281 | | |
| CD83 | 0.272 | | |
| PLD4 | 0.270 | | |
| CD79A | 0.269 | | |
| CD3E | 0.268 | | |
| CD1C | 0.267 | | |
| CD4 | 0.256 | | |
| BASP1 | 0.256 | | |

**Supplementary Table 5** | Gene association results corresponding to the second-ranked concept in EGFR predictions (TCGA-LUAD). Genes with Pearson's $r > 0.2$ are displayed.

| **Gene** | **Correlation** | **Gene** | **Correlation** |
|---|---|---|---|
| ACE2 | 0.413 | AIF1 | 0.218 |
| PROX1 | 0.406 | SNTN | 0.218 |
| PCSK2 | 0.398 | APCDD1 | 0.217 |
| GPX2 | 0.395 | CD1A | 0.209 |
| MALL | 0.381 | | |
| CAPN8 | 0.372 | | |
| CXCR4 | 0.359 | | |
| STEAP4 | 0.313 | | |
| TNFRSF17 | 0.307 | | |
| FCGR1A | 0.306 | | |
| ANPEP | 0.305 | | |
| ESR1 | 0.302 | | |
| VWA5A | 0.286 | | |
| TMC5 | 0.283 | | |
| EGFL7 | 0.283 | | |
| INMT | 0.281 | | |
| EGFR | 0.279 | | |
| PTPRC | 0.271 | | |
| CYP2A7 | 0.266 | | |
| MZB1 | 0.266 | | |
| CD83 | 0.265 | | |
| CXCL2 | 0.263 | | |
| FCER1A | 0.260 | | |
| DNAAF1 | 0.259 | | |
| CD1E | 0.256 | | |
| GPR183 | 0.256 | | |
| SEMA3C | 0.254 | | |
| SELL | 0.249 | | |
| LTBP2 | 0.245 | | |
| CCL5 | 0.245 | | |
| CD1C | 0.245 | | |
| CD14 | 0.244 | | |
| PLD4 | 0.243 | | |
| PDGFRA | 0.237 | | |
| KLRB1 | 0.234 | | |
| ADAM28 | 0.231 | | |
| CD79A | 0.228 | | |
| S100A1 | 0.225 | | |
| SOX2 | 0.221 | | |
| MYLK | 0.221 | | |

**Supplementary Table 6 | Prompt templates for CONCH** We used the same prompt templates as the original paper[63] for using CONCH model. We replace CLASSNAME with the actual name of the class.

```
CLASSNAME.
a photomicrograph showing CLASSNAME.
a photomicrograph of CLASSNAME.
an image of CLASSNAME.
an image showing CLASSNAME.
an example of CLASSNAME.
CLASSNAME is shown.
this is CLASSNAME.
there is CLASSNAME.
a histopathological image showing CLASSNAME.
a histopathological image of CLASSNAME.
a histopathological photograph of CLASSNAME.
a histopathological photograph showing CLASSNAME.
shows CLASSNAME.
presence of CLASSNAME.
CLASSNAME is present.
an H\&E stained image of CLASSNAME.
an H\&E stained image showing CLASSNAME.
an H\&E image showing CLASSNAME.
an H\&E image of CLASSNAME.
CLASSNAME, H\&E stain.
CLASSNAME, H\&E.
```

**Supplementary Table 7 | Class prompts for identifying tumor region.**

| Class | Class names |
|---|---|
| normal | non-tumor<br>normal tissue<br>non-cancerous tissue<br>benign |
| tumor | tumor tissue<br>cancerous tissue<br>malignant<br>cancer |
